# Chronic lung diseases are associated with gene expression programs favoring SARS-CoV-2 entry and severity

**DOI:** 10.1101/2020.10.20.347187

**Authors:** Linh T. Bui, Nichelle I. Winters, Mei-I Chung, Chitra Joseph, Austin J. Gutierrez, Arun C. Habermann, Taylor S. Adams, Jonas C. Schupp, Sergio Poli, Lance M. Peter, Chase J. Taylor, Jessica B. Blackburn, Bradley W. Richmond, Andrew G. Nicholson, Doris Rassl, William A. Wallace, Ivan O. Rosas, R. Gisli Jenkins, Naftali Kaminski, Jonathan A. Kropski, Nicholas E. Banovich, the Human Cell Atlas Lung Biological Network

## Abstract

Patients with chronic lung disease (CLD) have an increased risk for severe coronavirus disease-19 (COVID-19) and poor outcomes. Here, we analyzed the transcriptomes of 605,904 single cells isolated from healthy and CLD lungs to identify molecular characteristics of lung cells that may account for worse COVID-19 outcomes in patients with chronic lung diseases. We observed a similar cellular distribution and relative expression of SARS-CoV-2 entry factors in control and CLD lungs. CLD epithelial cells expressed higher levels of genes linked directly to the efficiency of viral replication and innate immune response. Additionally, we identified basal differences in inflammatory gene expression programs that highlight how CLD alters the inflammatory microenvironment encountered upon viral exposure to the peripheral lung. Our study indicates that CLD is accompanied by changes in cell-type-specific gene expression programs that prime the lung epithelium for and influence the innate and adaptive immune responses to SARS-CoV-2 infection.

## Introduction

In December 2019, a respiratory disease associated with a novel coronavirus emerged in Wuhan, China ^1–3^. The syndrome, now called COVID-19, was caused by severe acute respiratory syndrome coronavirus 2 (SARS-CoV-2) and has since rapidly spread worldwide ^4^. As of December 1, 2020, a total of over 63.4 million confirmed COVID-19 cases and more than 1.4 million deaths have been reported around the globe (https://coronavirus.jhu.edu/).

The clinical manifestations of SARS-CoV-2 infection range from asymptomatic to fulminant cases of acute respiratory distress syndrome (ARDS) and life-threatening multi-system organ failure. Development of ARDS in patients with SARS-CoV-2 dramatically increases the risk of ICU admission and death ^5–11^. Risk factors for severe SARS-CoV-2 include age, smoking status, ethnicity and male sex ^12–14^. Baseline comorbidities including hypertension, diabetes and obesity, increase SARS-CoV-2 susceptibility and severity ^9,15–19^. In addition, chronic lung disease (CLD) has been identified as a risk factor for hospitalization and mortality in patients with COVID-19 ^20–27^. Patients with chronic obstructive pulmonary disease (COPD) and interstitial lung disease (ILDs), especially Idiopathic Pulmonary Fibrosis (IPF), have a significantly higher COVID-19 mortality rate compared to patients without chronic lung disease ^28^. However, the molecular mechanisms underlying the increased risk of SARS-CoV-2 severity and mortality in patients with pre-existing lung diseases are not well understood.

To investigate the molecular basis of SARS-CoV-2 severity and mortality risk in CLD patients, we performed an integrated analysis of four lung single cell RNA-sequencing (scRNA-seq) datasets ^29–32^ in addition to unpublished data, together including 78 control and 132 CLD samples (n=31 COPD, 82 IPF and 19 other interstitial lung diseases). We found that CLD is associated with baseline changes in cell-type specific expression of genes related to viral replication and the immune response, as well as evidence of immune exhaustion and altered inflammatory gene expression. Together, these data provide a molecular framework underlying the increased risk of SARS-CoV-2 severity and poor outcomes in patients with certain pre-existing CLD.

## Results

### Expression profile of SARS-CoV-2 associated receptors and factors in the diseased lung

To determine why COVID-19 patients with CLD have a higher risk of severe infection and poorer outcomes, we performed an integrated analysis of the transcriptomes from 605,904 single cells derived from healthy donors (78 samples), COPD (31 samples), IPF (82 samples) and Non-IPF ILD (Other ILD, 19 samples) (Supplementary Table 1, 2). Using published cell type specific markers ^31,32^, we identified 38 distinct cell types in the dataset (Supplementary Fig. 1, Supplementary Table 3).

SARS-CoV-2 utilizes the host *ACE2,* and other putative factors such as *BSG, NRP1* and *HSPA5*, as entry receptor and *TMPRSS2*, *CTSL* or *FURIN* as priming proteases to facilitate cellular entry ^33–40^. Consistent with prior reports analyzing normal lung tissue ^33,34,41^, *ACE2* and *TMPRSS2* are expressed predominantly in epithelial cell types (Fig. 1a), while other putative SARS-CoV-2 entry receptors (*BSG*, *NRP1*, *HSPA5*) and priming proteases (*CTSL*, *FURIN*) have substantially more widespread expression in nearly all cell types (Supplementary Fig. 2). The number of *ACE2+* cells is highest in type 2 alveolar cells (AT2) and secretory cells, while *TMPRSS2* is widely expressed in all epithelial cell types. There were no significant differences in the proportion of *ACE2+* cells in any cell-type in diseased versus control groups (Fig. 1b). The proportion of *TMPRSS2+* AT2 cells are decreased in IPF lungs while *TMPRSS2+ SCGB3A2*+/*SCGB1A1*+ club cells are in significantly higher numbers in COPD and IPF patients compared to controls (Fig. 1c). The putative priming protease *CTSL* is expressed in more ciliated cells, proliferating macrophages, fibroblasts and pericytes isolated from Other-ILD samples (Supplementary Fig. 2).

**Fig. 1:**
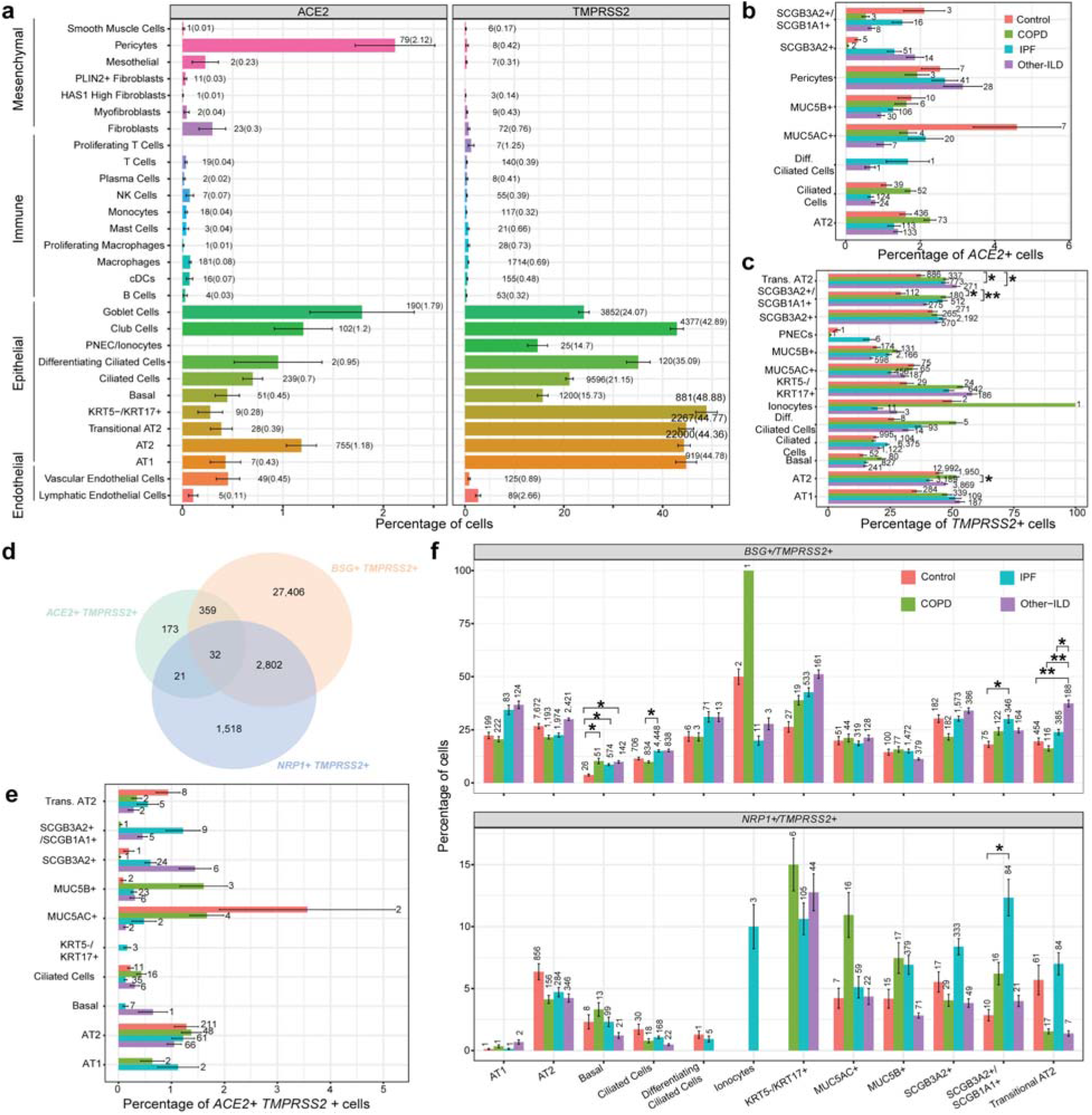
Percentage of cells expressing SARS-CoV-2 receptor genes in lung cell types in different diagnosis subgroups. (**a**) Percentage of cells expressing *ACE2* and *TMPRSS2* in all cell types. Total number of *ACE2*+ or *TMPRSS2*+ cells in the dataset (Percentage of *ACE2*+ or *TMPRSS2*+ cells). (**b-c**) Percentage of cells expressing *ACE2* and *TMPRSS2* in each diagnosis group, only cell types with at least 0.5% of cells expressing *ACE2* (**b**) and at least 10% of cells expressing *TMPRSS2* (**c**) are represented. (**d**) Venn diagram shows overlapping of cells co-expressing the proposed receptors (*ACE2*, *BSG* and *NRP1*) and the protease *TMPRSS2*. (**e-f**) Percentage of cells co-expressing receptors and *TMPRSS2* split by cell type and diagnosis group. Significant differences between diagnosis groups were calculated using Tukey_HSD test, p-value < 0.05: *, p-value < 0.01: **, p-value < 0.001: ***, p-value < 0.0001: ****.

Next, we compared the number of double positive cells, *i.e.* cells co-expressing a receptor and priming protease, in control and disease samples. Interestingly, a notable fraction of cells co-expresses all established and putative entry receptors (*ACE2*, *BSG*, *NRP1*, *HSPA5*) and proteases (*TMPRSS2*, *CTSL*, *FURIN*) (Fig. 1d, Supplementary Fig. 3). While the percentage of cells co-expressing *ACE2* and priming proteases (*TMPRSS2*, *CTSL*, *FURIN*) was similar across disease subtypes (Fig. 1e, Supplementary Fig. 3), there are significant differences in the number of cells co-expressing *BSG*, *NRP1*, *HSPA5* with a priming protease in disease samples in multiple cell types (Fig. 1f, Supplementary Fig. 3). Interestingly, the number of *BSG+/TMPRSS2+* and *NRP1+*/*TMPRSS2+* cells is significantly higher in the *SCGB1A1+/SCGB3A2+* club cells isolated from IPF samples.

To examine whether CLD patients express higher levels of SARS-CoV-2 receptors and priming proteases, we performed differential expression analysis of those genes in the disease versus control samples. The two major SARS-CoV-2 cellular entry factors, *ACE2* and *TMPRSS2*, have similar expression profiles in the disease and control samples. *ACE2* expression is highest in AT2 and Ciliated Cells, but there were no significant differences in *ACE2* expression in disease groups compared to control (Supplementary Fig. 4a). The putative alternative receptor *NRP1*, recently confirmed as another host entry factor for SARS-CoV-2^35^, is up-regulated in the COPD macrophages, but down-regulated in both IPF and Other-ILD macrophages (Supplementary Fig. 5). *TMPRSS2* expression is highest in AT2 cells, and is lower in the COPD samples (Supplementary Fig. 4b), similar to a recent publication demonstrating decreased *TMPRSS2* expression in severe COPD ^42^. Two alternative priming proteases (*CTSL* and *FURIN*) showed no significant differences in expression between control and disease samples (Supplementary Fig. 4e, f). However, the SARS-CoV-2 entry gene score (calculated on the average expression levels of all SARS-CoV-2 entry factors over a random control gene set) is significantly increased in the disease samples in many epithelial cell types, including AT1, AT2, Basal and Club cells, but not in *KRT5-/KRT17+* cells, an ECM-producing epithelial cell type enriched in the fibrotic lung ^31,32^, (Fig. 2b, Supplementary Fig. 6a,b). Together, these data suggest CLD is associated with only modest changes in expression of established SARS-CoV-2 entry factors, and an alternative mechanism likely is responsible for observed differences in outcome severity.

**Fig. 2:**
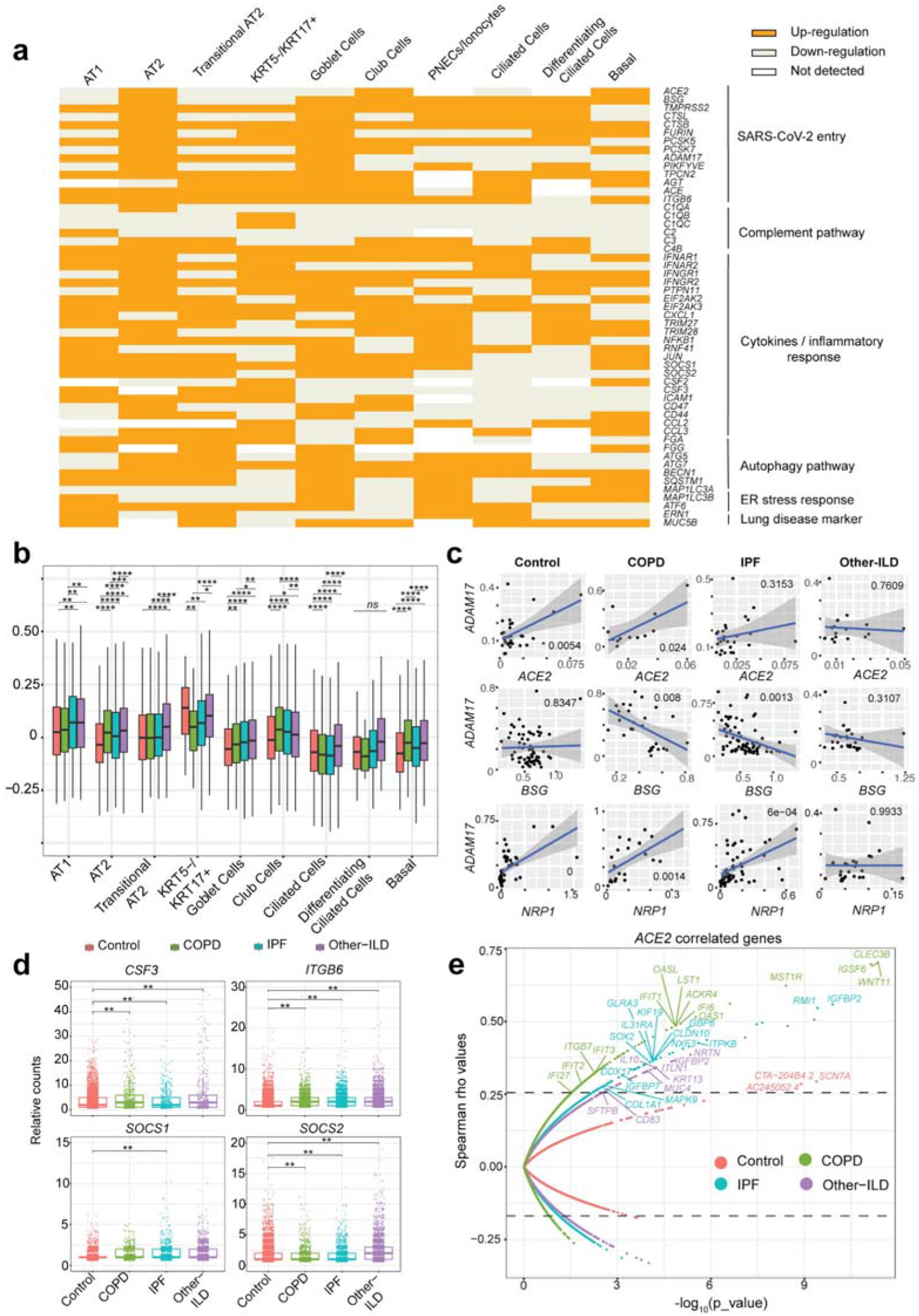
Expression profile of SARS-CoV-2 mediators and response genes in the epithelial cell population. (**a**) Binary heatmap with log_2_FC converted into a 1 (genes upregulated in the Disease samples) and 0 (genes downregulated in the Disease samples) shows the majority of the SARS-CoV-2 response genes are upregulated in the Disease samples compared to Control, the upregulation pattern in the disease samples were tested with an Exact binomial test, p-value = 9.909e-07. Up-regulation: genes up-regulated in Disease samples, Down-regulation: genes down-regulated in Disease samples, Not detected: gene expression was not detected in either of the two tested populations (Disease vs. Control). (**b**) SARS-CoV-2 entry module score in different cell types, SARS-CoV-2 mediators included *ACE2, BSG (CD147), NPR1, HSPA5 (GRP78), TMPRSS2, CTSL, ADAM17, FURIN*. The outliers were removed in this plot, please see Supplementary Fig. 6a with outliers included. *: p-value < 0.05, **: p-value < 0.01, ***: p-value < 0.001, ****: p-value < 0.0001, Tukey_HSD post-hoc test. (**c**) Significant gene expression correlation in AT2 cells between *ADAM17* and *ACE2*, *BSG* (*CD147*) and *NPR1* in COPD and IPF samples. (**d**) Boxplot shows differences in gene expression of selected Covid-19 response genes in the AT2 cell types, **: p_value_adj < 0.05 (negative binomial test, corrected for Age, Ethnicity, Smoking_status and Dataset). (**e**) Spearman gene correlation analysis identified genes correlated with *ACE2* expression in AT2 *ACE2+* cells in different diagnosis groups, p_value was adjusted using Benjamini-Hochberg corrections and dashed lines indicate the 99th percentile of Spearman rho values.

### Dysregulation of viral infection and innate immune response genes in disease epithelial cells

Given the relatively modest differences in SARS-CoV-2 entry factors in epithelial cells between diseased and control lungs (Supplementary Fig. 4, 5), we hypothesized that rather than greatly increased cellular susceptibility to SARS-CoV-2 infection, patients with CLD are predisposed to severe lung injury due to underlying differences in epithelial gene expression in key pathways mediating the antiviral response. Epithelial cells isolated from diseased lungs expressed higher levels of genes thought to directly impact viral replication (*PIKFYVE*, *TPCN2*) and the immune response to viral infection (Fig. 2a). The complement pathway genes *C3* and *C4B*, important components of the innate immune response and previously found to be elevated in SARS patients ^43^, are upregulated in diseased samples. Interestingly, key regulators of the host viral response including cytokine and inflammatory response genes (IFN type I and type II receptors, *SOCS1*/*2*, *CCL2, CSF3, TRIM28, EIF2AK2*) are also differentially expressed in diseased epithelial cells. Additionally, autophagy *(ATG5, ATG7, BECN1*) and ER stress response *(EIF2AK3, ATF6, ERN1*) genes which are well known to be aberrantly regulated in IPF lungs were also upregulated in epithelial cells in other types of chronic lung disease; these pathways are important for propagating viral infection and the host response ^44–46^.

In the distal lung, AT2 cells have been proposed to be the primary targets of SARS-CoV-2 ^34,41,47^. Thus, we examined the gene expression profile of diseased AT2 cells in more detail. As described above, AT2 cells in all disease subgroups have significantly higher SARS-CoV-2 entry gene scores than control cells (Fig. 2b). Additionally, genes involved in the complement pathway, immune response and autophagy pathway are upregulated in diseased AT2 cells (Fig. 2a). COPD and Other-ILD, but not IPF, AT2 cells express higher levels of *CSF3*, an important cytokine in the regulation of granulocytes (Fig. 2d). The epithelial integrin *ITGB6,* involved in wound healing and pathogenic fibrosis ^48^, and the suppressors of cytokine signaling-1 and −2 (*SOCS1, SOCS2*) are upregulated in all disease AT2 cells (Fig. 2d, Supplementary Fig. 6). Gene correlation analysis showed strong positive correlation between *ADAM17*, the main enzyme responsible for *ACE2* shedding, and *ACE2*, *NRP1* but negatively correlated with *BSG* and *HSPA5* in the COPD and IPF samples (Fig. 2c, Supplementary Fig. 7). Interestingly, *NRP1* expression is also positively correlated with the proteases *TMPRSS2* and *CTSL* in the AT2 cells isolated from IPF samples (Supplementary Fig. 7c). These data suggest that there are basal differences in the expression profiles of genes regulating viral infection and the immune response in diseased epithelial cells, specifically AT2 cells, and that this epithelial “priming” may contribute to COVID-19 severity and poor outcomes.

### Disease specific ACE2 correlated gene profiles in AT2 cells

Since *ACE2* is the best-established SARS-CoV-2 entry factor, we sought to identify *ACE2-* correlated genes in the *ACE2+* AT2 cells in different disease groups. Thus, identifying the immediate cellular environment SARS-CoV-2 encounters upon infecting a host. We performed Spearman correlation analysis with Benjamini-Hochberg adjusted p-values and identified distinct gene profiles significantly correlated with *ACE2* for each disease group (Fig. 2e). There were only three *ACE2* correlated genes in the Control samples with a cut-off of 99th percentile Spearman rho values and q-value less than 0.03, none of those genes are associated with the immune response. In the disease samples, we identified 671 genes (COPD: 429 genes, IPF: 140 genes and Other-ILD: 102 genes) with significant correlation to *ACE2* (99th percentile rho values, q-value less than 0.03) (Supplementary Table 7). Interestingly, many *ACE2-*correlated genes in the disease samples are associated with the innate and antiviral immune response. In the COPD samples, genes with strong correlation coefficients with *ACE2* include several interferon-induced genes (*IFI6, IFI27, IFIT1, IFIT2, IFIT3*), integrin beta 7 (*ITGB7*), a modulator of innate immune function (*OAS1*), the chemokine receptor *ACKR4* and a gene associated with West Nile viral infection (*OASL*). In the IPF samples, *ACE2* expression is strongly correlated with the interferon-induced gene *GBP6* (Guanylate Binding Protein Family Member 6), the interleukin receptor *IL31RA*, and *DDX17,* a gene downstream of the IFN pathway. In other ILD diseases (non-IPF related), the cytokine *IL10*, a member of the TGF-f] superfamily (*NRTN*), and an innate immune pathway component (*ITLN1*) are among the genes with high correlation with *ACE2*. Interestingly, *IFIT3* and *GBP4* (a member in the same gene family of *GBP6*) were shown to correlate with *ACE2* in asthma patients ^49^, while *OAS1* was among the top 50 genes with a significant correlation coefficient with *ACE2* in a previous study ^33^. The presence of immune-associated genes in these gene correlation profiles suggests that in patients with CLD, *ACE2+* AT2 cells are conditioned and primed to express these genes to cope with viral infection.

### Elevated ACE2 protein expression level in the small airway in IPF lungs

To further study the expression of the major SARS-CoV-2 entry factor *ACE2* in the fibrotic lungs, we examined the protein level of ACE2 in different lung regions. In agreement with the transcript quantification above and previous immunohistology analysis ^50^, we detected overall low expression level of ACE2 across all tissue types in both IPF (Fig. 3a-c) and control lung sections (Supplementary Fig. 8a). Notably, semi-quantitative evaluation of ACE2 expression scoring showed elevated ACE2 expression in all IPF sections compared to control, reaching statistical significance in the IPF small airway sections (Fig. 3a, g), suggesting that while overall ACE2 expression is low, there is a regional concentration of ACE2+ cells within the distal IPF lung that may promote a more severe localized viral response. Upregulation of the epithelial integrin alpha-V beta-6 (αvβ6) plays an important role in enhanced fibrosis in response to lung injury ^51^, and enhances TGFβ activation which can suppress type I interferon responses from alveolar macrophages increasing susceptibility to viral infection ^52^. We detected a significant increase of αvβ6 integrin expression in all lung sections isolated from IPF patients (Fig. 3d-f, h). While there was additional positive staining in the peripheral lung, αvβ6 expression is highest in the AT2 epithelial cells in the IPF samples compared to overall low expression level in the normal lung sections (Supplementary Fig. 8b), mirroring the expression data of *ITGB6* described above (Fig. 2d).

**Fig. 3:**
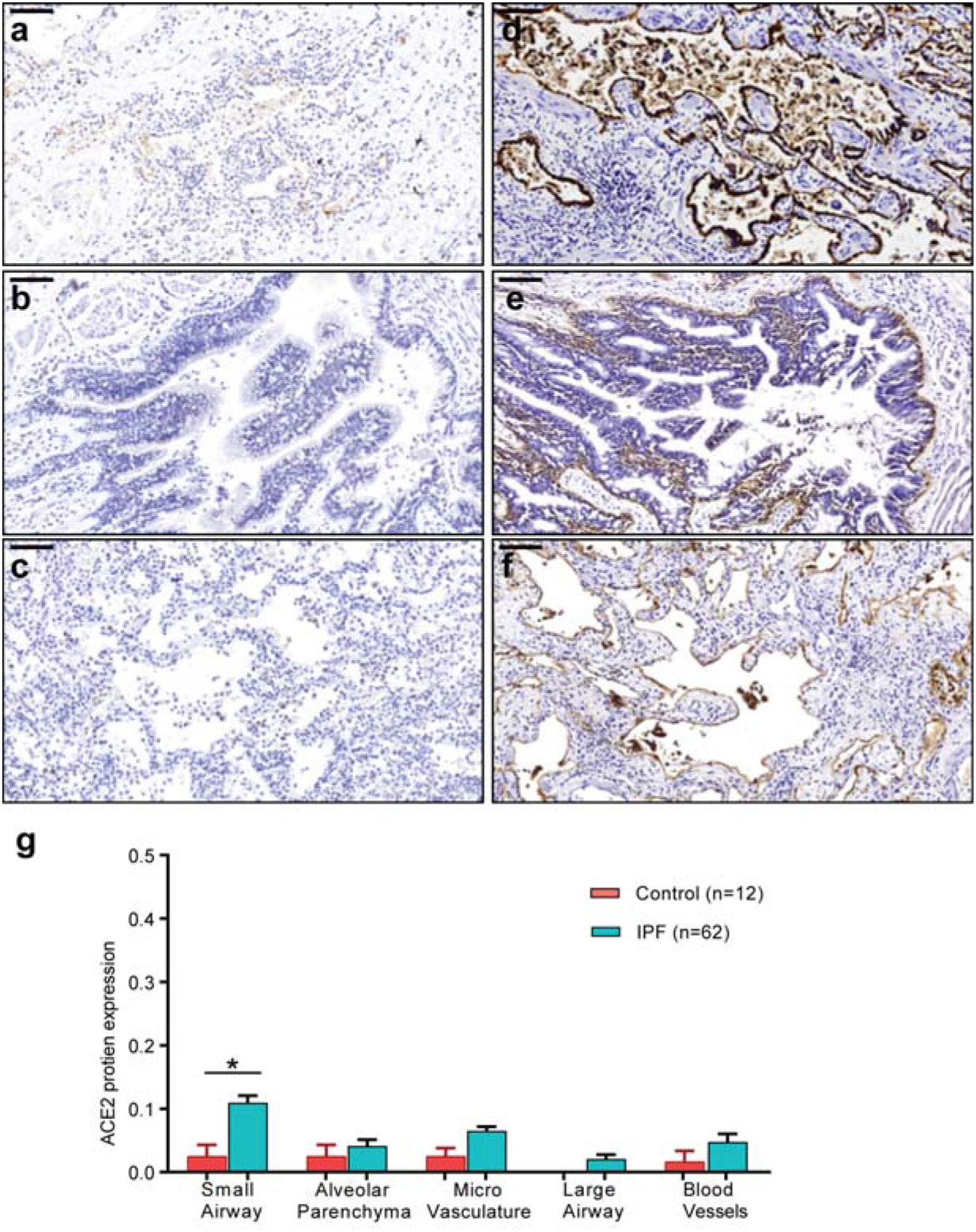
ACE2 and ITGB6 protein expression in IPF lung sections. (**a-c**) IPF lung sections stained for ACE2: (**a**) small airway, (**b**) large airway and (**c**) lung parenchyma. (**d-f**) IPF lung sections stained for αvβ6: (**d**) small airway, (**e**) large airway and (**f**) lung parenchyma. (**g**) semi-quantitative evaluation of ACE2 scoring among control and IPF sections (both the percentage of staining and staining intensity of ACE2 expression;0-Negative; 1-0–⍰10%; 2-11–⍰25%; 3-⍰26%). Significant differences between IPF and control were calculated using Tukey HSD test, p-value < 0.05 *. Scale bar = 100μm.

### Baseline differences in inflammatory response programs in chronic lung disease

Recent publications have suggested that immune dysregulation, including sustained cytokine production and hyper-inflammation, is associated with SARS-CoV-2 severity ^53–56^. We performed in-depth examination of the immune cell population to determine whether pre-existing immune dysregulation in chronic lung disease patients could contribute to SARS-CoV-2 severity and mortality. We analyzed a total of 418,891 cells from 13 immune cell types (Supplementary Table 3, Supplementary Fig. 1) and found significant increases in the proportion of CD4, CD8, cDCs and NK cells in the disease groups, most notably in COPD samples (Fig. 4a). Moreover, many immune cells isolated from disease samples expressed higher levels of genes in the interferon pathway, IL6-related cytokine pathway (*IL6ST*, *TGFB1, AREG*) and especially major histocompatibility complex (MHC) class II genes (HLA type II genes) (Fig. 4b). Expression of HLA type II genes increased across all disease groups but especially in the Other-ILD samples, compared to controls (Fig. 4d). Type I IFN response (IFNa score) is slightly elevated in the diseased macrophages and pDCs (Supplementary Fig. 9a). IL6-associated tocilizumab responsive genes (IL6 score) are expressed at a higher level in the disease groups IPF and Other-ILD, but lower in the COPD samples (Supplementary Fig. 9b). Previous studies demonstrated elevated exhaustion levels in CD8 T cells in severely affected COVID-19 patients ^57,58^. Notably, diseased immune cells, especially CD8 T cells and NK cells, have higher expression levels of cytotoxicity and exhaustion genes compared to controls (Fig. 4e, f). These perturbations in the T Cell population of diseased lungs may diminish the host immune response to viral infection, leading to a higher risk of severe disease and poor outcomes in response to SARS-CoV-2 infection.

**Fig. 4:**
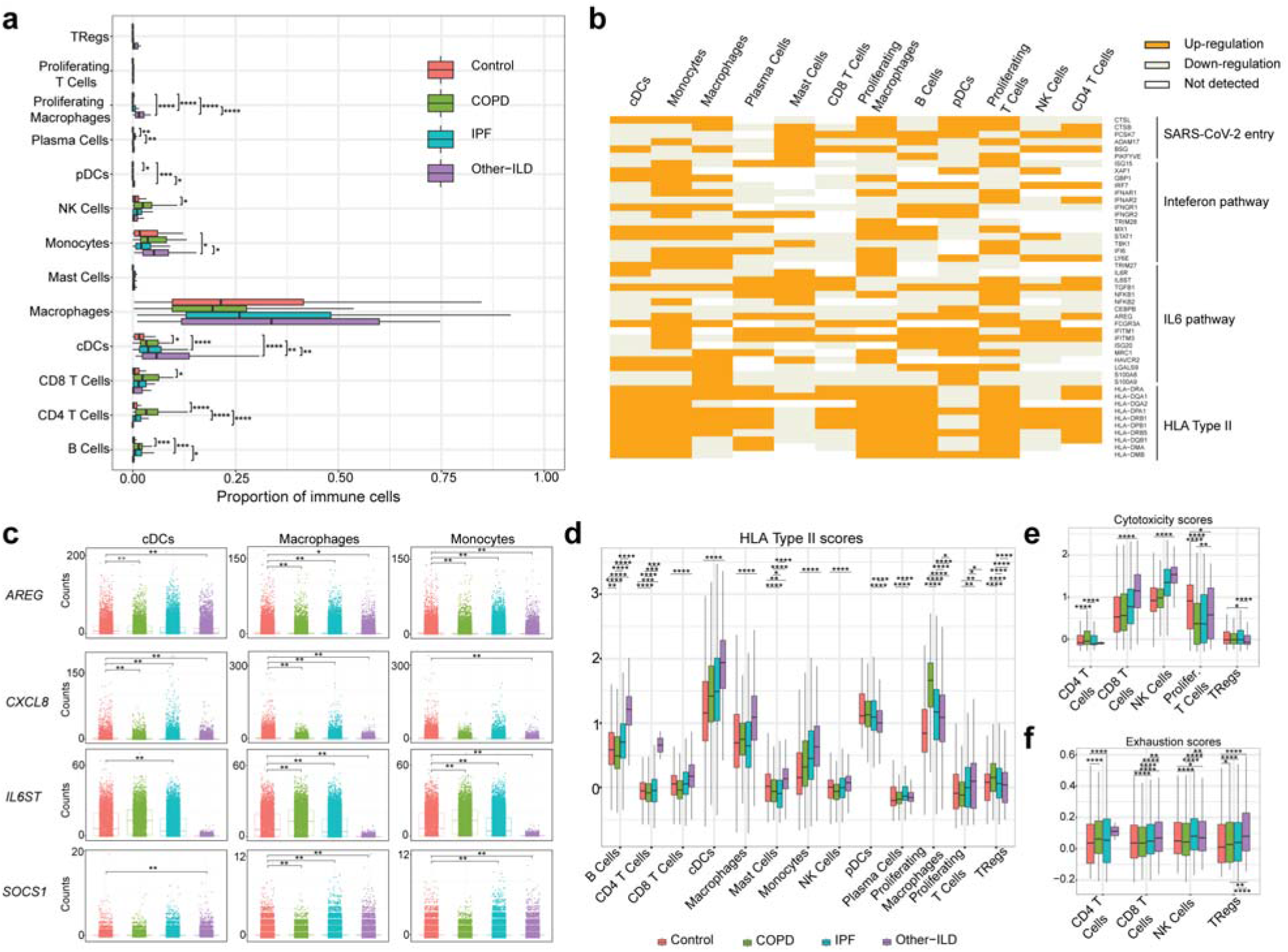
Analysis of SARS-CoV-2 candidate immune response genes in immune cells. (**a**) quantification of cell types as a percent of all immune cells in control versus diseased lungs. (**b**) Binary heatmap shows the majority of the SARS-CoV-2 response genes are upregulated in the Disease samples compared to Control; p-value = 8.298e-08 (Exact binomial test); Up-regulation: genes being up-regulated in Disease samples, Down-regulation: genes being down-regulated in Disease samples, Not detected: gene expression was not detected in either of the two tested populations (Disease vs. Control). (**c**) Differential expression analysis for SARS-CoV-2 immune candidate genes in cDCs, Macrophages and Monocytes. *: p_adjusted_value < 0.1, **: p_adjusted_value < 0.05, p_adjusted_value was Bonferroni adjusted from Seurat FindMarkers differential expression analysis using a negative binomial test and corrected for Age, Ethnicity, Smoking_status and Dataset. (**d**) Compared to the healthy control samples, HLA type II score is higher in all disease groups (especially Other-ILD). (**e-f**) In the T cell population, cytotoxicity scores (**e**) and exhaustion scores (**f**) are higher in the disease samples than in control samples. (**a**), (**d**), (**e**) and (**f**): Boxes: interquartile range, *: p-value < 0.05, **: p-value < 0.01, ***: p-value < 0.001, ****: p-value < 0.0001, Tukey_HSD post-hoc test. See Supplementary Fig. 10 for plots with outliers included for (**d**), (**e**), (**f**).

To further investigate differences in immune cell type-specific gene expression profiles, we examined expression levels of genes associated with viral infection in disease versus control samples. Amphiregulin (AREG), a ligand for epidermal growth factor receptor (EGFR), is known to have essential roles in wound repair and inflammation resolution; furthermore, upregulation of *AREG* is associated with viral infections of the lung ^59^. In COVID-19 patients, *AREG* is significantly upregulated in PBMCs ^60^, monocytes, CD4 T Cells, NK cells, neutrophils and DCs ^56^, suggesting that upregulation of *AREG* may be an attempt to ameliorate the severe injury induced by SARS-CoV-2 infection. We observed reduced expression of *AREG* in the cDCs and macrophages, but not in the monocytes, in the disease samples (Fig. 4c, Supplementary Table 6, Supplementary Fig. 10). *SOCS1*, a suppressor of cytokine signaling, was shown to reduce the type I IFN antiviral response in bronchial epithelial cells after influenza infection ^61,62^. Expression of the chemokine *CXCL8* and the IL6 co-receptor *IL6ST* was elevated in COVID-19 patients ^63,64^. In our study, *CXCL8* expression is lower in the disease samples in cDCs, macrophages and monocytes while *SOCS1* expression is elevated in disease samples in these cell types (Fig. 4c, Supplementary Table 6). Interestingly, *IL6ST* expression level is elevated significantly in COPD but reduced dramatically in Other-ILD samples (average two-fold down-regulation compared to Control samples in all three cell types) (Fig. 4c, Supplementary Table 6). These basal differences in inflammatory gene expression programs highlight how chronic lung disease alters the inflammatory microenvironment encountered upon viral exposure to the peripheral lung.

## Discussion

The COVID-19 pandemic, caused by the SARS-CoV-2 virus, has affected tens of millions of individuals around the globe in just the first nine months of 2020. Patients with CLD have an increased risk for severe SARS-CoV-2 infection: COPD patients have a five-fold increased risk of severe COVID-19 ^23,24,65–67^ and ILD patients have up to a four-fold increased odds of death from COVID-19 ^28,68^. Here, we performed an integrated transcriptomic analysis of scRNA-seq data from healthy and CLD patients to identify potential molecular causative factors determining SARS-CoV-2 severity. To summarize the results: (1) *ACE2* and *TMPRSS2* are expressed predominantly in epithelial cells and there are no significant differences in the number of *ACE2*+ cells in all cell types in disease compared to control samples; (2) a viral entry score including multiple entry factors is increased in cells isolated from diseased lungs; (3) diseased epithelial cells exhibit pre-existing dysregulation of genes involved in viral infection and the immune response; (4) a unique *ACE2* correlated gene profiles for each diagnosis group included anti-viral and immune regulatory genes; (5) ACE2 protein levels are elevated in the IPF small airway sections; (6) there are baseline differences in the cellular immune population in disease compared to control samples.

Similar to other coronaviruses, SARS-CoV-2 utilizes cellular receptors (*ACE2* and putatively, *BSG, NRP1* and *HSPA5* gene products) and priming proteases (*TMPRSS2, CTSL, FURIN*), for viral entry. These factors are expressed predominantly in the upper and lower airways, with *ACE2* being expressed highly in nasal goblet and ciliated cells and in a subset of AT2 cells and the absorptive enterocytes in the gut ^33,34,41,47,50^. We observed a similar expression pattern of *ACE2* in our dataset, with AT2 cells having the highest number *ACE2+* cells. To our knowledge, publications investigating baseline expression of these SARS-CoV-2 entry factors in lung disease have been limited to asthma and COPD with variable results. For example, studies in asthma patients showed elevated expression of *ACE2, TMPRSS2* and *FURIN* in patients with severe but not mild-moderate asthma ^49,69^. Leung *et al.* performed bulk RNAseq and immunohistochemical staining on bronchial epithelial cells and showed significantly elevated expression levels of *ACE2* and ACE2 protein in the small airways of COPD patients compared to control ^70^. Another study on bronchoscopically isolated tissue showed no relationship between disease status (mild to moderate asthma or COPD) on the expression levels of all SARS-CoV-2 entry factors ^42^. Our study utilized scRNAseq technology to study gene expression at a very granular level and did not identify increased *ACE2* expression at the single cell level in CLD, including COPD. However, in the IPF lung, there was a regional concentration of ACE2+ cells in the small airways upon immunohistochemical examination (Fig. 3a, g), similar to the findings of Leung *et al ^70^*. While the overall frequency of *ACE2*+ cells and the ACE2 expression level may be low, changes in the proportional cellular makeup of the diseased lung epithelium may lead to a proportionate increase in *ACE2*+ “infectable” cells into the distal lung. Importantly, IPF lungs exhibit abnormal expansion of epithelial cell programs, specifically the presence of more proximal specific cell types in the distal lungs ^31,32^. Thus, our data along with previously published studies together suggest that while overall differences in *ACE2* expression and other entry factors may be minimal in CLD, the localization of susceptible cells in the distal lung may promote disease pathogenesis and severity.

A balanced immune response is crucial to viral clearance and avoidance of excessive injury to the host, as evidenced by poor outcomes related both to immunosuppression as well as hyperinflammation in COVID-19 patients^71^. COPD patients with severe COVID-19 had elevated serum levels of various inflammatory cytokines including IL-2R, IL-6, IL-8, IL-10 and TNF-α suggesting there may be global alterations in the immune response ^27^.We observed that COPD AT2 cells expressed elevated levels of immune-response related genes (*CSF3, ITGB6, SOCS1, SOCS2*). G-CSF (encoded by *CSF3)* is found at high levels in patients with severe COVID-19 and thought to play a role in the hyperinflammatory syndrome while *SOCS1* and *SOCS2* are part of a negative feedback system that regulates the response to cytokines ^72,73^. *ACE2* correlated genes in this cell population were enriched for regulators of the immune response (Fig. 2e), with several of these genes found to be upregulated in alveolosphere cultures infected with SARS-CoV-2 ^36^. In addition to alterations in the cytokine microenvironment, changes in cellular immune populations were also identified in the COPD samples, dysregulation of several genes in inflammatory pathways (*AREG, CCL3, CXCL8, SOCS1*), and high levels of cytotoxic and exhaustion-related genes in CD4 and CD8 T Cells from COPD lungs. Expression of cytotoxic and exhaustion genes was increased compared to controls but similar in IPF and Other-ILD immune cell types. Together, our data suggest that the immune microenvironment at both the molecular and cellular level in the fibrotic and COPD lung is dysregulated and may promote severe infection and poor outcomes in COVID-19.

One limitation of our study is that we focus mainly on the peripheral regions of the lungs, and did not analyze cells in the upper airways or trachea. It is possible that there are significant differences in SARS-CoV-2 entry gene expression between disease and control samples in the more proximal regions of the lungs. Our study is also limited to the expression profiles of patients with CLD without SARS-CoV-2 infection, as the collection of samples from patients who are both infected with SARS-CoV-2 and have chronic lung disease are difficult to collect at present. Nevertheless, our study highlights that dysregulation of genes related to viral replication and innate immune response in the epithelial cells, and basal differences in the inflammatory gene expression programs are key factors leading to an increased risk of COVID-19 severity and poorer outcomes in patients with CLD.

## Methods

The code for genomic analyses in this paper is available at https://github.com/tgen/banovichlab/

### scRNA-seq samples

scRNA-seq data were obtained from published data with samples in the “VUMC/TGen” dataset from Habermann *et. al.* (2020) (GEO accession GSE135893), samples in the “Yale/BWH” dataset came from Adams *et. al.* (2020) (GEO accession number GSE136831), samples in the “Pittsburg” dataset from Morse *et. al.* (2019) (GEO accession GSE128033) and samples in the “Northwestern” dataset from Reyfman *et al.* (2019) (GEO accession GSE122960) (Supplementary Table 1, 2). Additionally, there are 39 unpublished scRNA-seq samples in the “VUMC/TGen” dataset that were collected under Vanderbilt IRB #’s 060165, 171657 and Western IRB # 20181836.

### scRNA-seq data processing

Seurat v3.1.5 package ^74^ was used to process the scRNA-seq data. Specifically, for the Pittsburgh, Northwestern datasets and 39 unpublished samples from the VUMC/TGen, the CellRanger (10X Genomics) output files were read into Seurat using the function *Read10X*, the remaining datasets were already in Seurat format and were loaded using the function *readRDS*. To eliminate low-quality/dying cells or empty droplets, we removed any cells containing fewer than 1,000 genes or more than 25% mitochondrial genes. Due to the large size of the joint dataset, we performed *SCTransform^75^* for normalization and scaling of each dataset separately, split into four major cell populations using known markers: *EPCAM*+ (Epithelial), *PECAM1*+ *PTPRC* - (Endothelial), *PTPRC* + (Immune) and *EPCAM-PECAM-PTPRC-* (Mesenchymal). Each population from the four datasets was then merged together to generate four merged Seurat objects (Endothelial, Epithelial, Immune and Mesenchymal). Next, each object was SCTransformed with “dataset” being used as batch_var to eliminate batch effects between datasets. Cell type annotation was manually performed on each object using known cell-type specific markers (Supplementary Fig. 1, ^32^). For each cell population, cell type annotation was performed at four levels, ranging from the most general to more granular annotation. After removing doublet cells, all four population datasets were merged to generate the final ILD object containing a total of 605,904 cells and 38 distinct cell types (Supplementary Table 2, Supplementary Fig. 1).

### Integrated analysis of joint dataset

To calculate the percentage of single positive or double positive cells for *ACE2* and other cofactors, we counted the number of cells with >0 transcripts of corresponding genes. For double positive, cells have >0 transcripts of both genes of interest. Tukey Honest Significant Difference (Tukey_HSD) statistical test from the R package *rstatix* with a confidence level of 0.95 was used to test statistical dependence of cells expressing the SARS-CoV-2 mediators among chronic disease subsets.

To assess the expression profile of SARS-CoV-2 mediators (*ACE2*, *BSG*, *NRP1, HSPA5)*, the corresponding proteases (*TMPRSS2*, *CTSL*, *FURIN*) and other candidate genes involved in SARS-CoV-2 infection in different chronic lung disease subset (COPD, IPF or Other-ILD), we ran the function *FindMarkers* in Seurat package using the negative binomial test. To account for batch effects, we used the parameter “*latent_vars*” to incorporate the four variables (Age, Ethnicity, Smoking status and Dataset) into the negative binomial model. For the binary heatmap, the differential expression analysis was performed between the Disease (including all chronic disease subset) and Control samples. Then, log_2_ fold-change was converted into 0 (downregulated in the disease samples) or 1 (upregulated in the disease samples) regardless of the Bonferroni adjusted p-values. Heatmaps were generated from the adjusted log_2_FC values using the *heatmap.2* function of the *gplots* R package ^76^, and exact binomial statistics tests were performed to test whether the selected genes are upregulated in the disease samples across cell types. For the boxplots, count numbers of selected genes were plotted using the *geom_boxplot* and *geom_jitter* function of the *ggplot2* R package ^77^. Significant differences in gene expression in the boxplots were the Bonferroni adjusted p-values calculated from the FindMarkers function between Control and Disease groups (COPD, IPF, Other ILD) using the fitted negative binomial model and latent_vars parameters as described above.

### Gene module score

To calculate the combined expression of genes, we used the *AddModuleScore* in Seurat v3.1.5 package. SARS-CoV-2 entry gene scores were calculated on SARS-CoV-2 receptors and mediators: *ACE2, BSG (CD147), NRP1, HSPA5(GRP78), TMPRSS2, CTSL, FURIN* and *ADAM17*. HLA type II score includes *HLA-DRA, HLA-DQA1, HLA-DQA2, HLA-DPA1, HLA-DRB1, HLA-DPB1, HLA-DQB2, HLA-DRB5, HLA-DQB1, HLA-DMA, HLA-DMB*. IFN score includes *ISG15, IFI44, IFI27, CXCL10, RSAD2, IFIT1, IFI44L, CCL8, XAF1, GBP1, IRF7, CEACAM1*. IL6 scores were calculated on six tocilizumab responsive genes: *ARID5A, SOCS3, PIM1, BCL3, BATF, MYC* that are associated with the IL-6 pathway ^56^. Cytotoxicity associated genes include *PRF1, GZMH, IFNG, NKG7, KLRG1, PRF1 ^56^* and *GNLY, GZMB, GZMK ^78^*. Exhaustion gene set: *LAG3, TIGIT, PDCD1, CTLA4, HAVCR2, TOX ^58^* and *PRDM1, MAF ^56^*. Significant differences between different groups were calculated using the Tukey_HSD statistic test in the R package *rstatix* with a confidence level of 0.95 (Supplementary Table 5).

### Gene correlation analysis

To identify genes that correlate with *ACE2* in the AT2 *ACE2+* cells, we performed Spearman correlation coefficient analysis on the log-transformed and normalized data using the function *cor.test* in the R stats v3.6.1 package with Benjamini-Hochberg corrections for p_adjusted values.

### Immunohistochemistry of ACE2 and anti-αvβ6 integrin

Formalin-fixed paraffin-embedded histological sections of human lung were cut at 5-microns and dewaxed in xylene prior to rehydration in decreasing concentrations of ethanol. The tissue samples were obtained after informed consent and local ethics approval (South East Scotland SAHSC Bioresource-reference number 06/S1101/41; Brompton Node samples - reference number 15/SC/0101; Papworth Node Samples; non-diseased controls-reference number (Q)GM030404 and Nottingham BRC samples-reference number 08/H0407/1). IHC staining was performed using the Novocastra Novolink™ Polymer Detection Systems kit (Code: RE7280-K, Leica, Biosystems, Newcastle, UK) as previously described ^79^. Heat-induced citrate antigen retrieval (pH 6.0) and pepsin antigen retrieval was performed for Rabbit monoclonal ACE2 (ab108252, EPR4435(2) Abcam, UK) and the anti-αvβ6 integrin antibody (6.2A1; Biogen, Cambridge, MA, USA), respectively. Rabbit monoclonal ACE2 (1:400) and anti-αvβ6 integrin (1:3000) was diluted in Leica antibody diluent (RE AR9352, Leica, Biosystems, UK) and incubated with the sections overnight at 4°C. Novolink DAB substrate buffer plus was used as the chromogen and the slides were counterstained with Novolink haematoxylin for 6 min, dehydrated and cover slipped. A negative control without the application of the primary antibody, and was also used to ensure staining was only related to the presence of the antibody.

The immunohistochemically stained slides were scanned using a ScanScope XT Slide Scanner (Leica Aperio Technologies, Vista, CA, USA) under 20x objective magnification (0.5μm resolution) using Pannoramic Viewer (3DHISTECH Ltd Budapest, Hungary) slide viewing software. Both the percentage of staining and staining intensity of ACE2 expression in lung sections were individually assessed. For ACE2 quantification, the following scoring system of six high-power fields at X40. per tissue section were used: The coding was performed prior to scoring/analysis as: **0**-Negative; **1**-0–⍰10%; **2**-11–⍰25%; **3**-⍰26%. Statistical analyses were completed using GraphPad Prism 7.0 (GraphPad Software, San Diego, CA, USA). One-way analysis of variance was used for comparison of more than two data sets and significant differences between diagnosis groups were calculated using Tukey HSD test.

## Data availability

The majority of the data used in this manuscript is publicly available from published paper: GEO accession GSE135893 ^32^, GEO accession GSE136831^31^, GEO accession GSE128033 ^30^ and GEO accession GSE122960 ^29^. The unpublished data from VUMC/TGen (39 samples) are included in the supplementary data (Supplementary Table 4) as a count matrix format containing all the genes being used in the manuscript.

## Acknowledgments

This study was supported by the NIH/NHLBI R01HL145372 (NEB/JAK), the Department of Defense W81XWH1910415 (NEB/JAK), Doris Duke Charitable Foundation (JAK), T32HL094296 (NIW, JBB), the Department of Veterans Affairs IK2BX003841 (BWR), DoD W81XWH-19-1-0131 (JCS), R01HL127349 (NK), R01HL141852 (NK), U01HL145567 (NK), UH2 HL123886 (NK), and a generous gift from Three Lakes Partners to NK and IOR. The integrated data sets were funded by various sponsors as indicated in the original publications. RGJ is funded by and NIHR Research Professorship (RP-2017-08-ST2-014).

## Author contributions

L.T.B, N.I.W., M.-I.C, N.E.B and J.A.K conceived and designed the analysis. A.J.G., L.T.B., N.E.B., and J.A.K. performed quality checks, data integration, and computational analyses. L.T.B., N.E.B., and J.A.K. analyzed and interpreted scRNA-seq data. C.J. performed the immunohistology and semi-quantification analysis. L.T.B., N.I.W., M.-I.C, N.E.B., and J.A.K. wrote and revised the manuscript, with significant input from I.O.R., G.J., N.K. and the HCA Lung Biological Network. All authors read and approved the manuscript before submission.

## Competing interests

JAK has received advisory board fees from Boehringer Ingelheim, Inc, Janssen Pharmaceuticals, is on the scientific advisory board of APIE Therapeutics, and has research contracts with Genentech. In the last 36 months, NK reported personal fees from Biogen Idec, Boehringer Ingelheim, Third Rock, Samumed, Numedii, AstraZeneca, Life Max, Teravance, RohBar, and Pliant and Equity in Pliant; collaboration with MiRagen, AstraZeneca; Grant from Veracyte, all outside the submitted work. In addition, NK has a patent for New Therapies in Pulmonary Fibrosis, and Peripheral Blood Gene Expression licensed to Biotech. AGN has received advisory board fees from Boehringer Ingelheim, Galapagos, Medical Quantitative Image Analysis and personal fees for educational material from Up to Date and Boehringer Ingelheim. RGJ reports grants from AstraZeneca, grants from Biogen, personal fees from Boehringer Ingelheim, personal fees from Chiesi, personal fees Daewoong, personal fees from Galapagos, grants from Galecto, grants from GlaxoSmithKline, personal fees from Heptares, non-financial support from NuMedii, grants and personal fees from Pliant, personal fees from Promedior, non-financial support from Redx, personal fees from Roche, other from Action for Pulmonary Fibrosis, outside the submitted work.

## Notes

https://github.com/tgen/banovichlab/

### Summary of Updates

Addition of NPR1 gene expression analysis, revision of title and text

