## Supplementary Tables and Figures for "Chronic lung diseases are associated with gene expression programs favoring SARS-CoV-2 entry and severity"

**Online Data Supplement**

HCA Lung Biological Network Members

Supplementary Tables 1-7

Supplementary Figures 1-10

Supplementary references

HCA Lung Biological Network Members

Alexander V. Misharin

Division of Pulmonary and Critical Care Medicine, Northwestern University, Chicago, Illinois

Alexander M. Tsankov

Genetics and Genomic Sciences, Icahn School of Medicine at Mount Sinai, New York, NY USA

Avrum Spira

Department of Medicine, Boston University School of Medicine, Boston, MA, USA; Johnson & Johnson Innovation, Cambridge, MA, USA.

Pascal Barbry

Université Côte d’Azur, CNRS, IPMC, Sophia-Antipolis, 06560, France

Alvis Brazma

European Molecular Biology Laboratory, European Bioinformatics Institute (EMBL-EBI), Wellcome Trust Genome Campus, Hinxton, Cambridge, CB10 1SD, UK

Christos Samakovlis

SciLifeLab, Department of Molecular Biosciences, Stockholm University, Stockholm Sweden and Cardiopulmonary Institute, Justus Liebig University; Giessen Germany

Douglas P Shepherd

Center for Biological Physics and Department of Physics, Arizona State University, Tempe, AZ USA

Emma L Rawlins

Wellcome Trust/ CRUK Gurdon Institute and Department Physiology, Development and Neuroscience, University of Cambridge

Fabian J Theis

Institute of Computational Biology, Helmholtz Zentrum München and Departments of Mathematics and Life Sciences, Technical University Munich, Germany

Jennifer Griffonnet

Nice University-Affiliated Hospital : Pneumology department, Nice, 06002 France

Haeock Lee

Department of Biomedicine and Health Sciences, The Catholic University of Korea, Seoul, Korea

Herbert B Schiller

Comprehensive Pneumology Center (CPC) / Institute of Lung Biology and Disease (ILBD), Helmholtz Zentrum München, Member of the German Center for Lung Research (DZL), Munich, Germany

Paul Hofman

Laboratory of Clinical and Experimental Pathology, Pasteur Hospital, University Côte d'Azur, Nice, France / Hospital‐Related Biobank, Pasteur Hospital, University Côte d'Azur, Nice, France / FHU OncoAge, Pasteur Hospital, BP69, 06001 Nice cedex 01, France

Joseph E Powell

Garvan-Weizmann Centre for Cellular Genomics, Garvan Institute of Medical Research, Sydney, NSW, Australia; UNSW Cellular Genomics Futures Institute, University of New South Wales, Sydney, NSW, Australia

Joachim L Schultze

Department for Genomics & Immunoregulation, LIMES-Institute, University of Bonn, 53115 Bonn, Germany

PRECISE Platform for Single Cell Genomics & Epigenomics, Germany Center for Neurodegenerative Diseases and University of Bonn, Bonn, Germany

Jeffrey Whitsett

Cincinnati Children’s Hospital Medical Center, Cincinnati, Ohio, USA

Jiyeon Choi

Division of Cancer Epidemiology and Genetics, National Cancer Institute, Bethesda, MD, 20892, USA.

Joakim Lundeberg

SciLifeLab, Department of Gene Technology, KTH Royal Institute of Technology

Jonathan A Kropski

Division of Allergy, Pulmonary and Critical Care Medicine, Department of Medicine, Vanderbilt University Medical Center, Nashville, TN; Department of Veterans Affairs Medical Center, Nashville, TN; Department of Cell and Developmental Biology, Vanderbilt University, Nashville, TN

Jose Ordovas-Montanes

Broad Institute of MIT and Harvard, Cambridge, MA 02142, USA; Program in Immunology, Harvard Medical School, Boston, MA 02115, USA; Division of Gastroenterology, Hepatology, and Nutrition, Boston Children's Hospital, Boston, MA 02115, USA; Harvard Stem Cell Institute, Cambridge, MA 02138, USA.

Jayaraj Rajagopal

Harvard Stem Cell Institute, Cambridge, Massachusetts; Center for Regenerative Medicine, Massachusetts General Hospital, Boston, Massachusetts, USA.

Kerstin B Meyer

Cellular Genetics Programme, Wellcome Sanger Institute, Wellcome Genome Campus, Hinxton, Cambridge CB10 1SA, United Kingdom.

Mark A Krasnow

Department of Biochemistry and Wall Center for Pulmonary Vascular Disease, Stanford University, Stanford, CA 94305, USA.

Kourosh Saeb‐Parsy

Department of Surgery, University of Cambridge and NIHR Cambridge Biomedical Research Centre, UK

Kun Zhang

UCSD Department of Bioengineering, 9500 Gilman Drive, MC0412, PFBH402, La Jolla, CA 92093, USA.

Robert Lafyatis

Division of Rheumatology, Department of Medicine, University of Pittsburgh Medical Center, Pittsburgh, PA, USA.

Sylvie Leroy

Université Côte d'Azur, CHU de Nice, FHU OncoAge, Department of Pulmonary Medicine and Allergology, Nice, France ; CNRS UMR 7275 - Institut de Pharmacologie Moléculaire et Cellulaire, Sophia Antipolis, France

Muzlifah Haniffa

Wellcome Sanger Institute, Wellcome Genome Campus, Hinxton, Cambridge CB10 1SA, UK; Biosciences Institute, Faculty of Medical Sciences, Newcastle University, Newcastle upon Tyne NE2 4HH, UK; Department of Dermatology and NIHR Newcastle Biomedical Research Centre, Newcastle Hospitals NHS Foundation Trust, Newcastle upon Tyne NE2 4LP, UK.

### Muzlifah Haniffa

Institute of Cellular Medicine, Newcastle University, Newcastle upon Tyne, U.K.

Department of Dermatology and Newcastle NIHR Biomedical Research Centre, Royal Victoria Infirmary, The Newcastle upon Tyne Hospitals NHS Foundation Trust, Newcastle upon Tyne, U.K.

Martijn C Nawijn

Department of Pathology and Medical Biology, University of Groningen, GRIAC Research Institute, University Medical Center Groningen, the Netherlands

Marko Z Nikolić

UCL Respiratory, Division of Medicine, University College London, London, UK

[Maarten van den Berge](https://pubmed.ncbi.nlm.nih.gov/?term=van+den+Berge+M&cauthor_id=31281062)

Department of Pulmonary Diseases, University Medical Center Groningen, and Groningen Research Institute for Asthma and COPD, University of Groningen, NL-9700-RB Groningen, Netherlands

Malte Kuhnemund

Cartana AB, Nobels vag 16, 17165 Stockholm, Sweden

[Charles-Hugo Marquette](https://pubmed.ncbi.nlm.nih.gov/?term=Marquette+CH&cauthor_id=30813420)

Team 4, IRCAN, FHU OncoAge, University Côte d'Azur, CNRS, INSERM, 06107 Nice CEDEX 02, France..

Department of Pneumology and Oncology, CHU Nice, FHU OncoAge, University Côte d'Azur, 06100 Nice, France..

Michael Von Papen

Comma Soft AG, Bonn, Germany

Naftali Kaminski

Pulmonary, Critical Care and Sleep Medicine, Yale University School of Medicine

Nicholas Banovich

Translational Genomics Research Institute

Oliver Eickelberg

Orit Rosenblatt-Rosen

Klarman Cell Observatory, Broad Institute of MIT and Harvard, Cambridge, MA 02142, USA

Paul A Reyfman

Northwestern University Feinberg School of Medicine, Division of Pulmonary and Critical Care Medicine

Dana Pe’er

Computational and Systems Biology Program, Sloan Kettering Institute, Memorial Sloan Kettering Cancer Center, New York, New York, USA

Peter Horvath

Biological Research Centre of the Hungarian Academy of Sciences, Temesvári krt. 62., 6726 Szeged Hungary.

Institute for Molecular Medicine Finland (FIMM), University of Helsinki, Tukholmankatu.

Purushothama Rao Tata

Department of Cell Biology, Regeneration Next Initiative, Duke University School of Medicine, Durham, NC, USA, 27710

Aviv Regev

Broad Institute of MIT and Harvard, Cambridge, MA 02142

Howard Hughes Medical Institute

Department of Biology, MIT, Cambridge, MA 02140

Genentech, 1 DNA Way, South San Francisco, CA 94080

Mauricio Rojas

Division of Pulmonary, Allergy and Critical Care Medicine, University of Pittsburgh

Max A Seibold

Department of Pediatrics; Center for Genes, Environment, and Health; National Jewish Health; Denver, CO 80206

Alex K Shalek

Ragon Institute of MGH, MIT, and Harvard, Cambridge, MA, USA; Institute for Medical Engineering and Science (IMES), Koch Institute for Integrative Cancer Research, and Department of Chemistry, Massachusetts Institute of Technology, Cambridge, MA, USA; Broad Institute of MIT and Harvard, Cambridge, MA, USA

Jason R Spence

Department of Internal Medicine, Gastroenterology, University of Michigan Medical School, Ann Arbor, MI 48109, USA; Department of Cell and Developmental Biology, University of Michigan Medical School, Ann Arbor, MI 48109, USA; Department of Biomedical Engineering, University of Michigan College of Engineering, Ann Arbor, MI 48109, USA.

Sarah A Teichmann

Cellular Genetics Programme, Wellcome Sanger Institute, Wellcome Genome Campus, Hinxton, Cambridge CB10 1SA, United Kingdom.

Dept Physics/Cavendish Laboratory, University of Cambridge, JJ Thompson Ave, Cambridge CB3 0EH, United Kingdom.

Stephen Quake

Chan Zuckerberg Biohub

Thu Elizabeth Duong

Department of Pediatrics, Respiratory Medicine, University of California, San Diego, USA

Tommaso Biancalani

Broad Institute of MIT and Harvard, Cambridge, MA 02142

Tushar Desai

Department of Medicine and Institute for Stem Cell Biology and Regenerative Medicine, Stanford University School of Medicine, Stanford, CA 94116

Xin Sun

Department of Pediatrics, University of California, San Diego, San Diego, CA 92093, USA.

Laboratory of Genetics, University of Wisconsin–Madison, Madison, WI 53706, USA.

Laure Emmanuelle Zaragosi

Université Côte d’Azur, CNRS, IPMC, Sophia-Antipolis, 06560, France

**SUPPLEMENTARY TABLES**

**Supplementary Table 1**: Number of samples used in the study

| **Dataset** | **Control** | **COPD** | **IPF** | **Other ILD** |
| --- | --- | --- | --- | --- |
| Northwestern | 9 | 0 | 4 | 4 |
| Pittsburg | 9 | 0 | 8 | 0 |
| VUMC/TGen | 22 | 7 | 25 | 15 |
| Yale/BWH | 38 | 24 | 45 | 0 |
| **Total** | **78** | **31** | **82** | **19** |

**Supplementary Table 2**: Demographics of lung donors used for scRNA-seq

|  | **Control** | **COPD** | **IPF** | **Other-ILD** |
| --- | --- | --- | --- | --- |
|  | n=78 | n=31 | n=82 | n=19 |
| Age | 44.23 [17-80] | 62 [55-73] | 64.87 [43-78] | 54.95 [37-70] |
| Ever_smoke | 29 (37.2%) | 28 (90.3%) | 52 (63.4%) | 8 (42%) |
| Gender (Male) | 44 (56.4%) | 19 (61.3%) | 59 (71.9%) | 10 (52.6%) |
| Ethnicity |  |  |  |  |
| European | 54 (69.2%) | 31 (100%) | 60 (73.2%) | 12 (63.2%) |
| African | 8 (10.2%) | - | 3 (3.7%) | 3 (15.8%) |
| Hispanic | 1 (1.3%) | - | 4 (4.9%) | - |
| Asian | 2 (2.6%) | - | 1 (1.2%) | - |
| Other/Unknown | 13 (16.7%) | - | 14 (17%) | 4 (21%) |

**Supplementary Table 3**: Total number of each cell type per diagnosis group

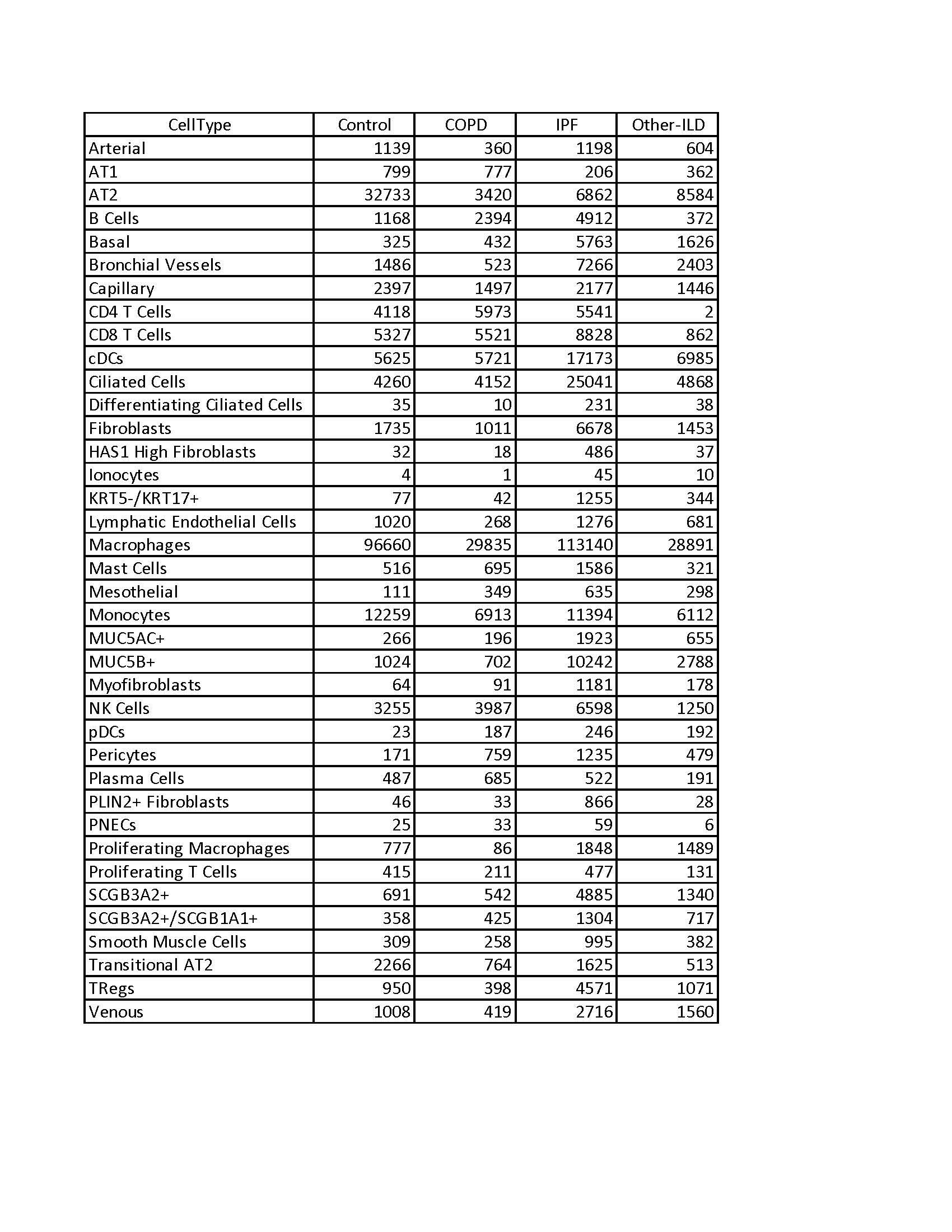

**Supplementary Table 4**: Count data matrix for unpublished data containing all genes being used in the manuscript (.csv file)

**Supplementary Table 5**: Excel sheets for all Tukey_HSD statistical tests (Excel file)

**Supplementary Table 6**: Differential expression analysis for genes used in Figure 2 and Figure 3 (Zip file)

**Supplementary Table 7**: Spearman correlation coefficient analysis for *ACE2* in AT2 cells (Excel file)

**SUPPLEMENTARY FIGURES**

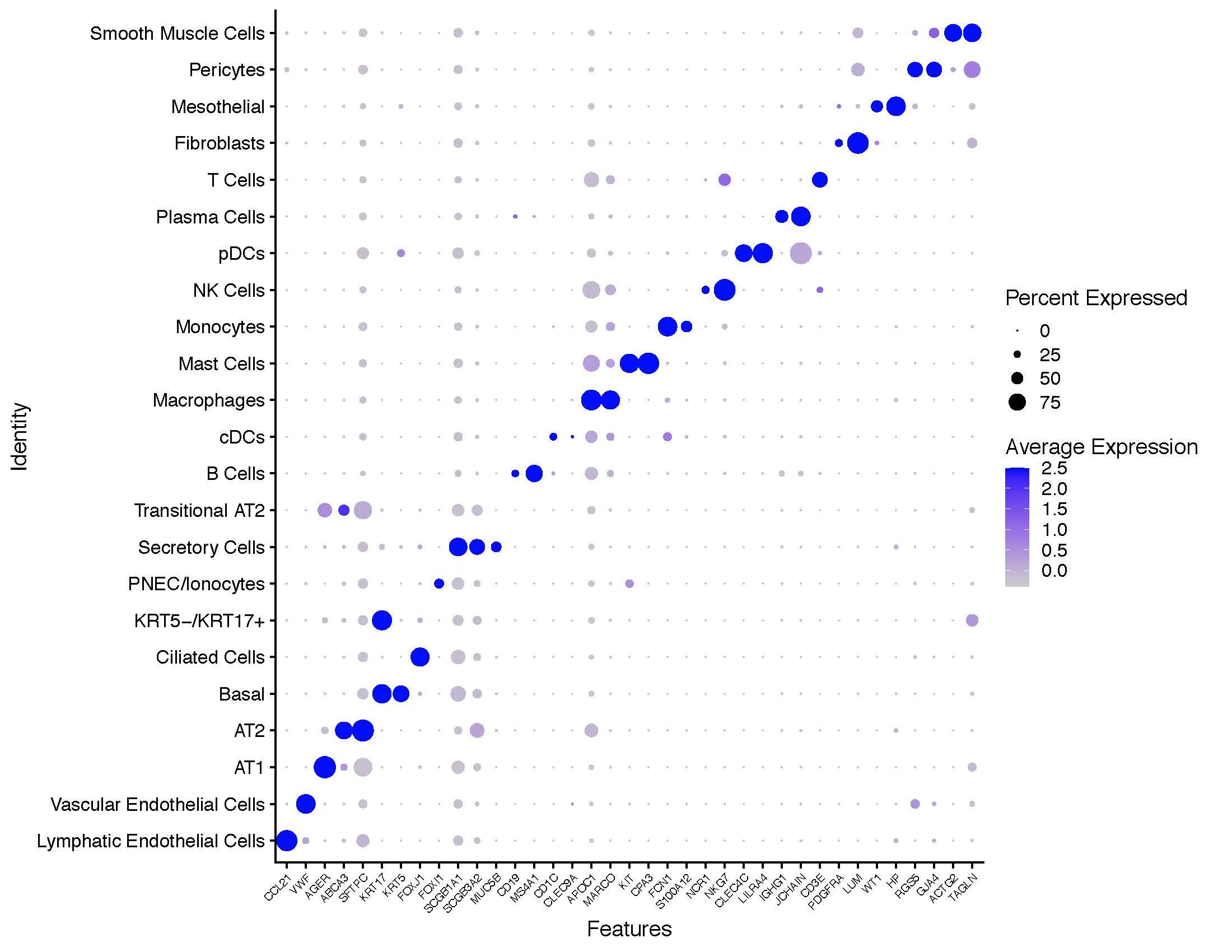

**Supplementary Fig 1**: Dotplot shows markers used for cell type annotation

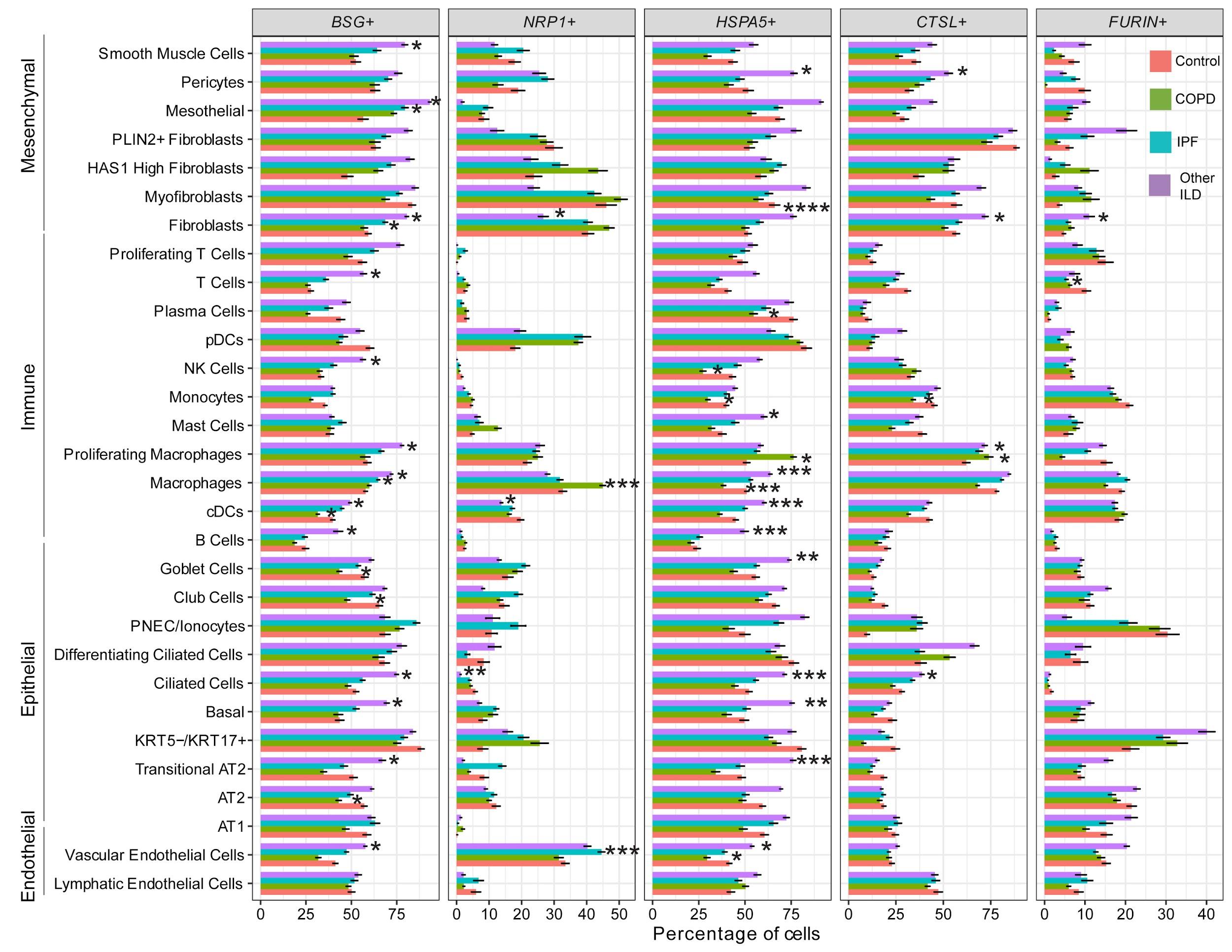

**Supplementary Fig 2**: Percentage of single positive cells for *BSG* (*CD147*), *NRP1,* *HSPA5* (*GRP78*), *CTSL* and *FURIN* in different chronic lung disease groups per cell type. Tukey_HSD post-hoc statistical test showed significant differences between each diagnosis subgroup and control samples, p-value < 0.05: *, p-value < 0.01: **, p-value < 0.001: ***, p-value < 0.0001: ****

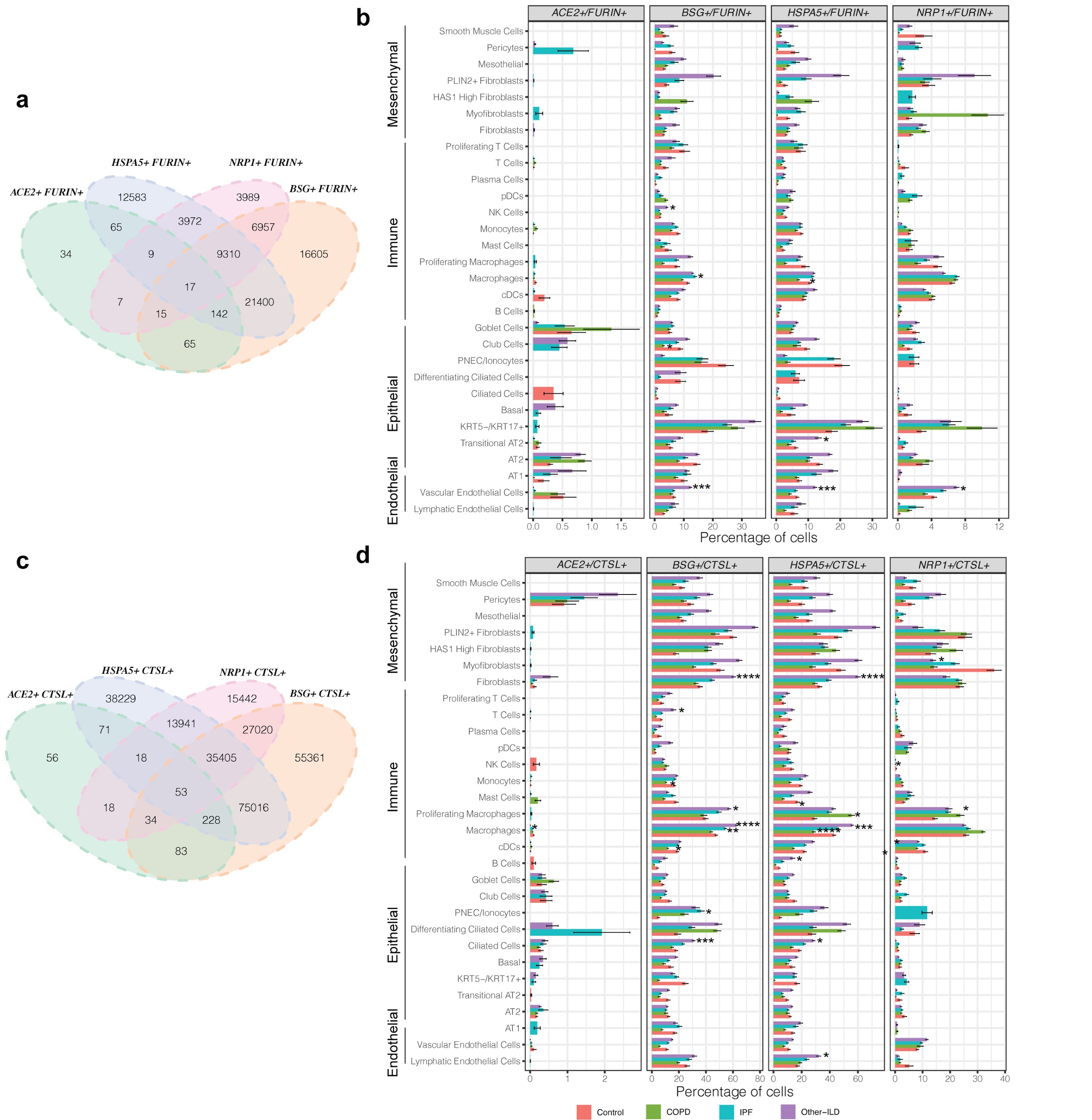

**Supplementary Fig 3**: Percentage of double positive cells: (**a-b**) *ACE2*+ *FURIN*+, *BSG*+ *FURIN*+, *HSPA5*+ *FURIN*+, *NRP1+ FURIN+* and (**c-d**) *ACE2*+ *CTSL*+, *BSG*+ *CTSL*+, *HSPA5*+ *CTSL*+, *NRP1+ CTSL+* in different chronic lung disease groups per cell type. Tukey_HSD post-hoc statistical test, p-value < 0.05: *, p-value < 0.01: **, p-value < 0.001: ***, p-value < 0.0001: ****

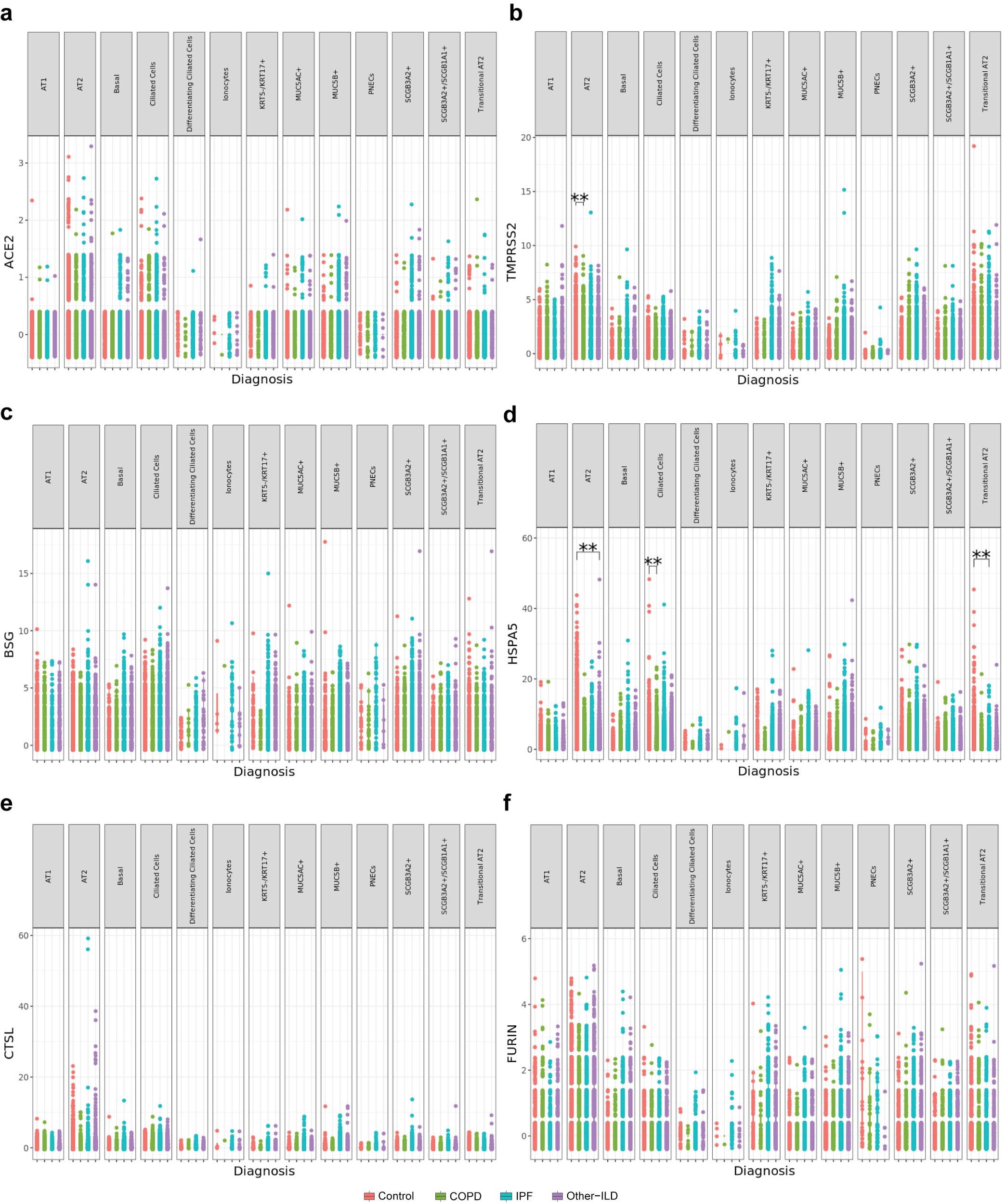

**Supplementary Fig 4**: Gene expression analysis for SARS-CoV-2 entry mediators in the epithelial cell population. (**a**) *ACE2*. (**b**) *TMPRSS2*. (**c**) *BSG* (*CD147*). (**d**) *HSPA5* (*GRP78*). (**e**) *CTSL*. (**f**) *FURIN*. **: Bonferroni adjusted p_value < 0.05 (Negative binomial test, corrected for Age, Ethnicity, Smoking_status and Dataset)

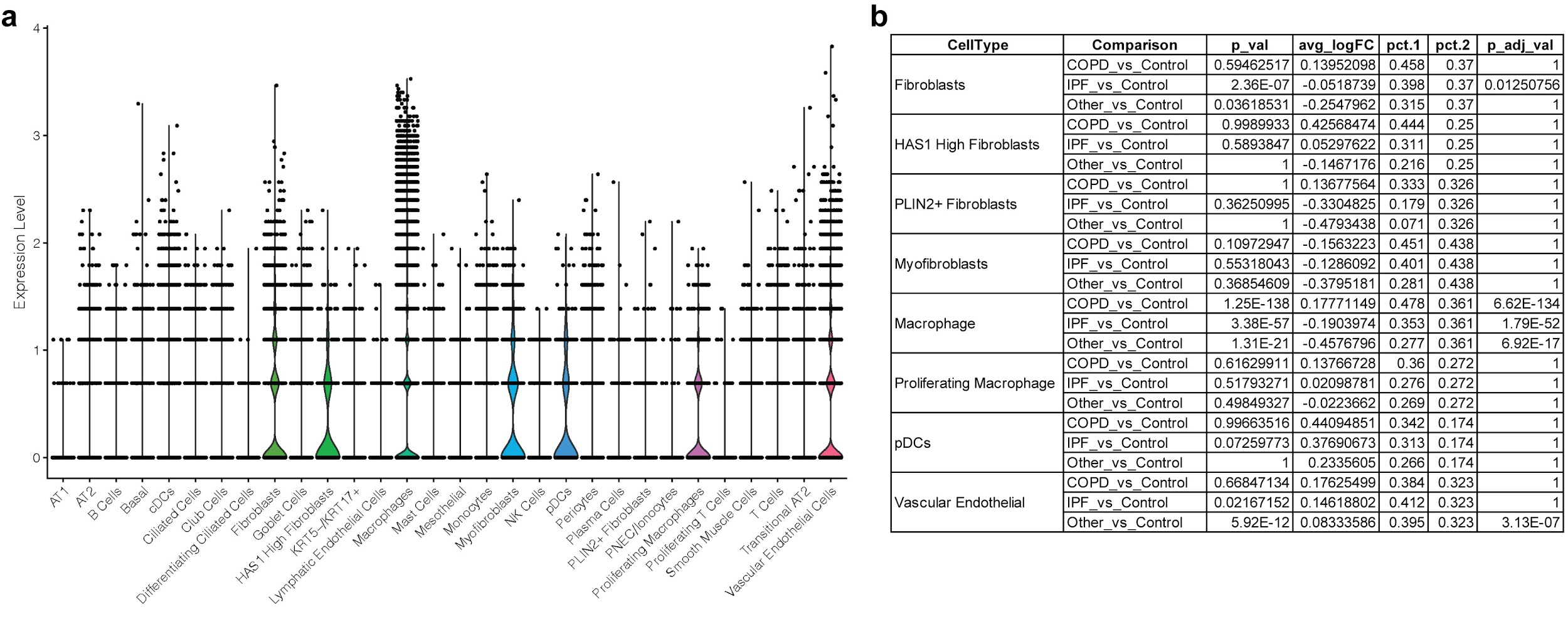

**Supplementary Fig 5**: Expression of the putative SARS-CoV-2 receptor, *NRP1*, in the data set. (**a**) Violin plot shows high expression of *NRP1* in some cell types. (**b**) Differential expression analysis of *NRP1* in different disease groups vs. control samples for the cell types having high *NRP1* expression in (**a**). The differential expression analysis was performed using the Seurat FindMarkers function and p_adj_val was Bonferroni adjusted (as described in the Methods section).

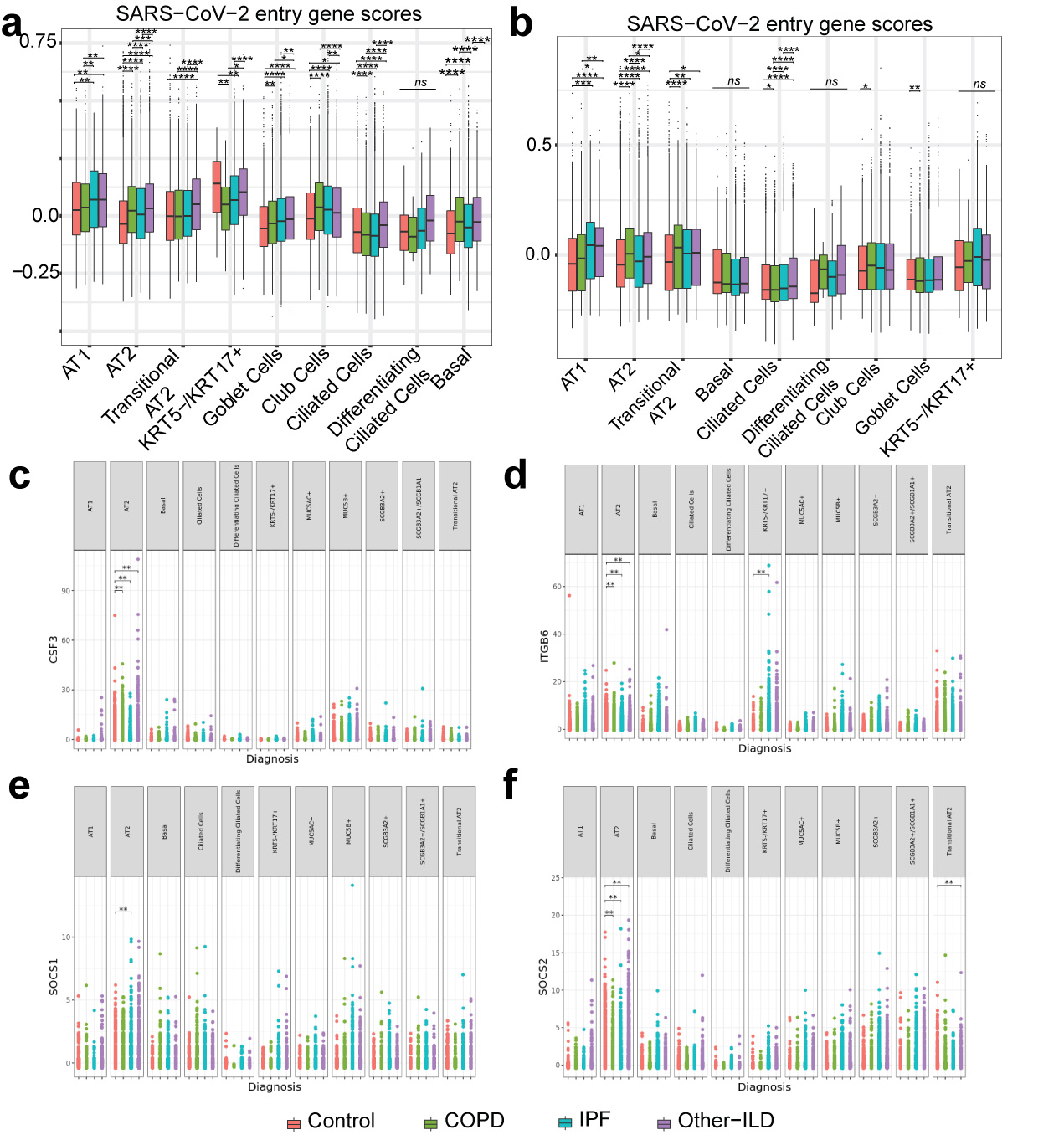

**Supplementary Fig 6**: SARS-Cov-2 entry genes scores with all outliers included. (**a**) Gene set 1 (*ACE2, BSG, HSPA5, TMPRSS2, CTSL, FURIN, ADAM17*). (**b**) Gene set 2 (*ACE2, TMPRSS2, CTSL, FURIN, ADAM17*). (**c-f**) differential expression of Covid-19 response genes in all epithelial cells, a full plot of Fig 2d. In (**a**) and (**b**): Tukey_HSD post-hoc statistical test, p-value < 0.05: *, p-value < 0.01: **, p-value < 0.001: ***, p-value < 0.0001: ****. In (**c**), (**d**), (**e**), (**f**): **: Bonferroni adjusted p_value < 0.05 (negative binomial test, corrected for Age, Ethnicity, Smoking_status and Dataset)

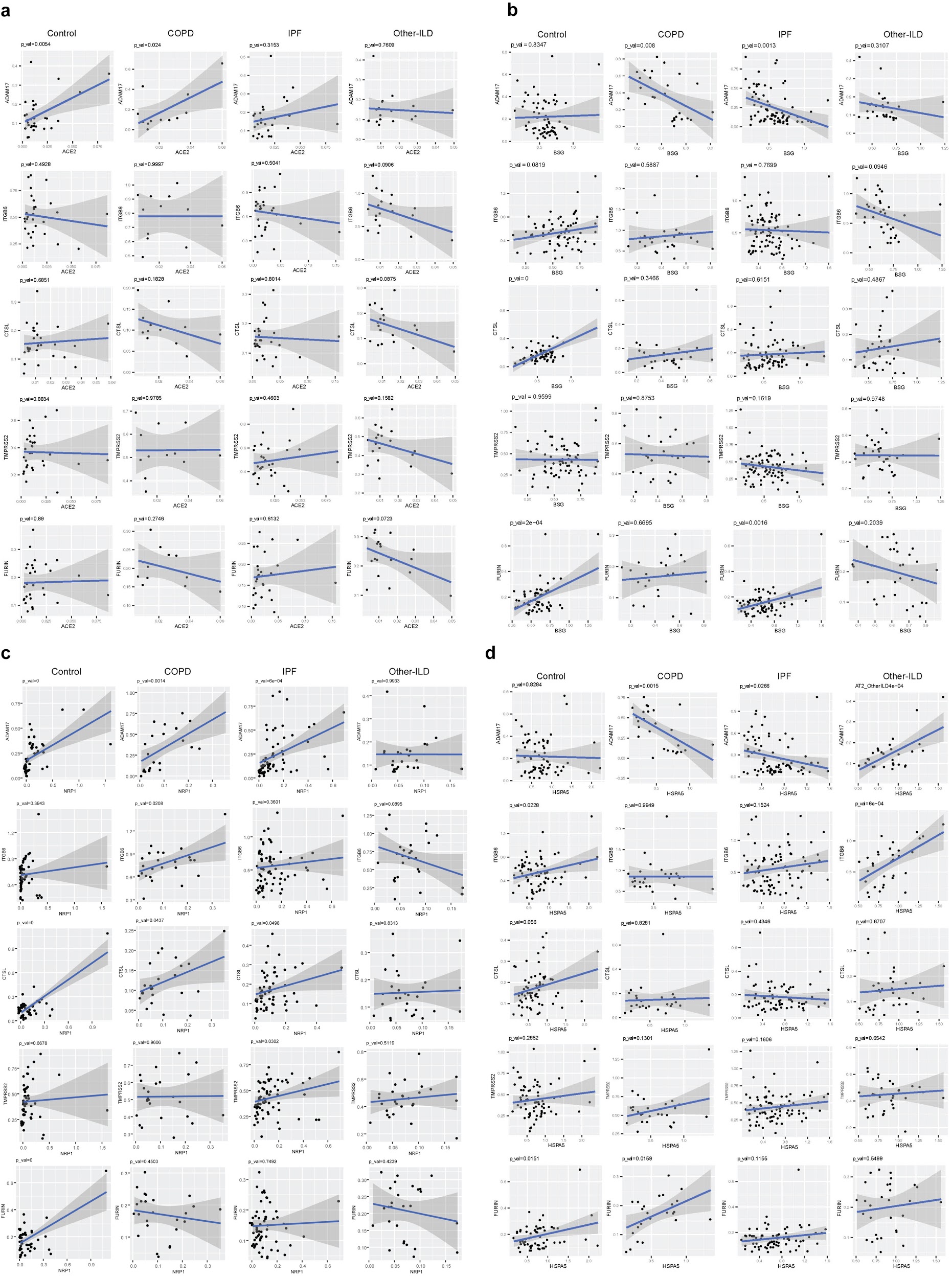

**Supplementary Fig 7**: Gene expression correlation between SARS-CoV-2 mediators and proteases in the AT2 cell types. (**a**) Gene correlation with *ACE2*. (**b**) Gene correlation with *BSG* (*CD147*). (**c**) Gene correlation with *NRP1*. (**d**) Gene correlation with *HSPA5* (*GRP78*)

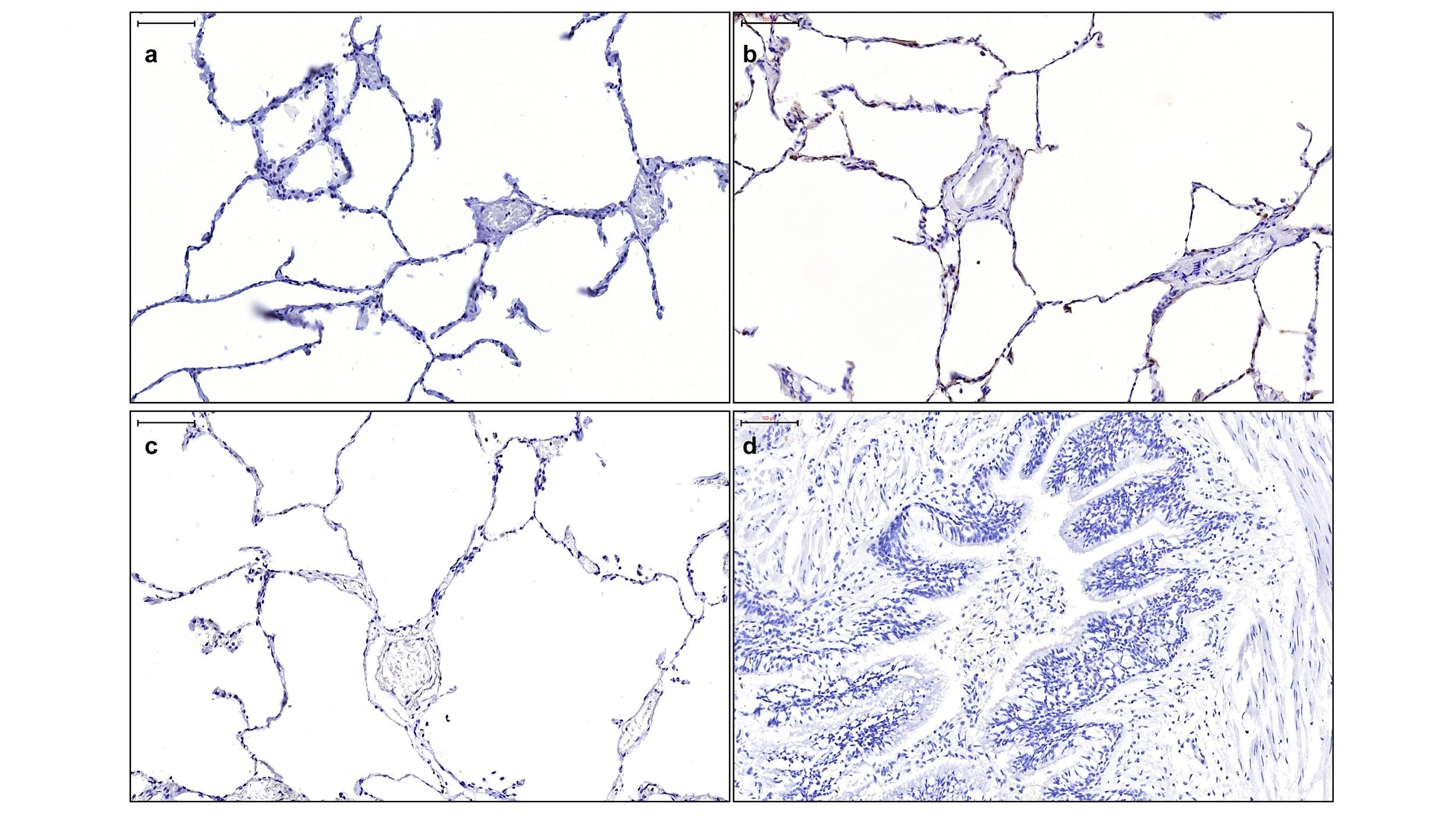

**Supplementary Fig 8**: Representative sections of control lung sections are shown with each immunomarker. (**a**) ACE2 staining of the alveolar parenchyma. (**b**) αvβ6 staining in the alveolar parenchyma. (**c-d**) Secondary antibody controls for control and IPF section; Scale bars=100 µm.

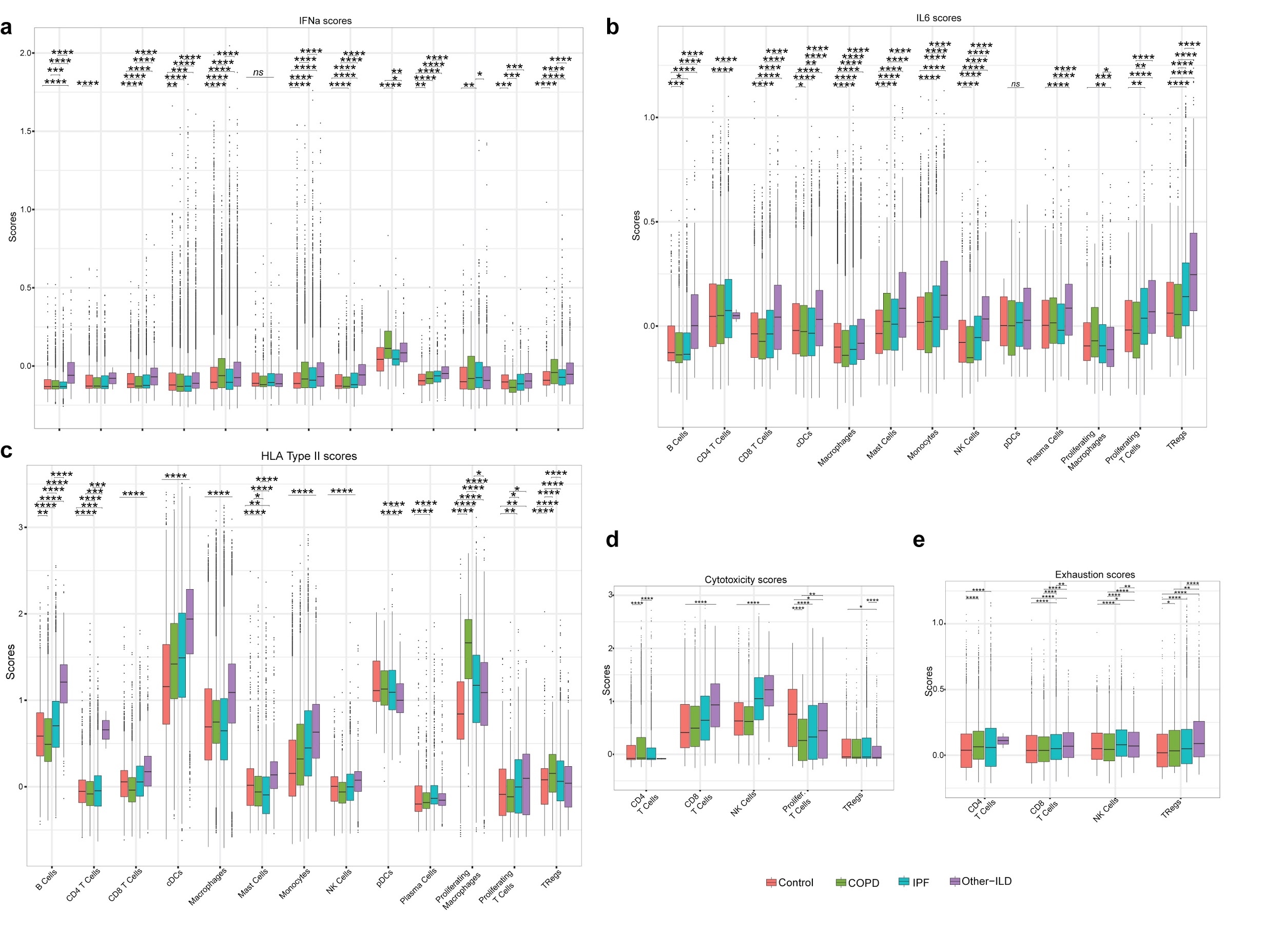

**Supplementary Fig 9**: Differences in average expression of IFNa (**a**), IL6 (**b**) and HLA type II (**c**) genes in the immune population in different disease groups. (**d-e**) Cytotoxic scores and exhaustion scores in the T cell population in different disease groups. (**b**), (**d**), (**e**) are the same plots as in Fig. 3d-f, but included outliers. Tukey_HSD post-hoc statistical test, p-value < 0.05: *, p-value < 0.01: **, p-value < 0.001: ***, p-value < 0.0001: ****

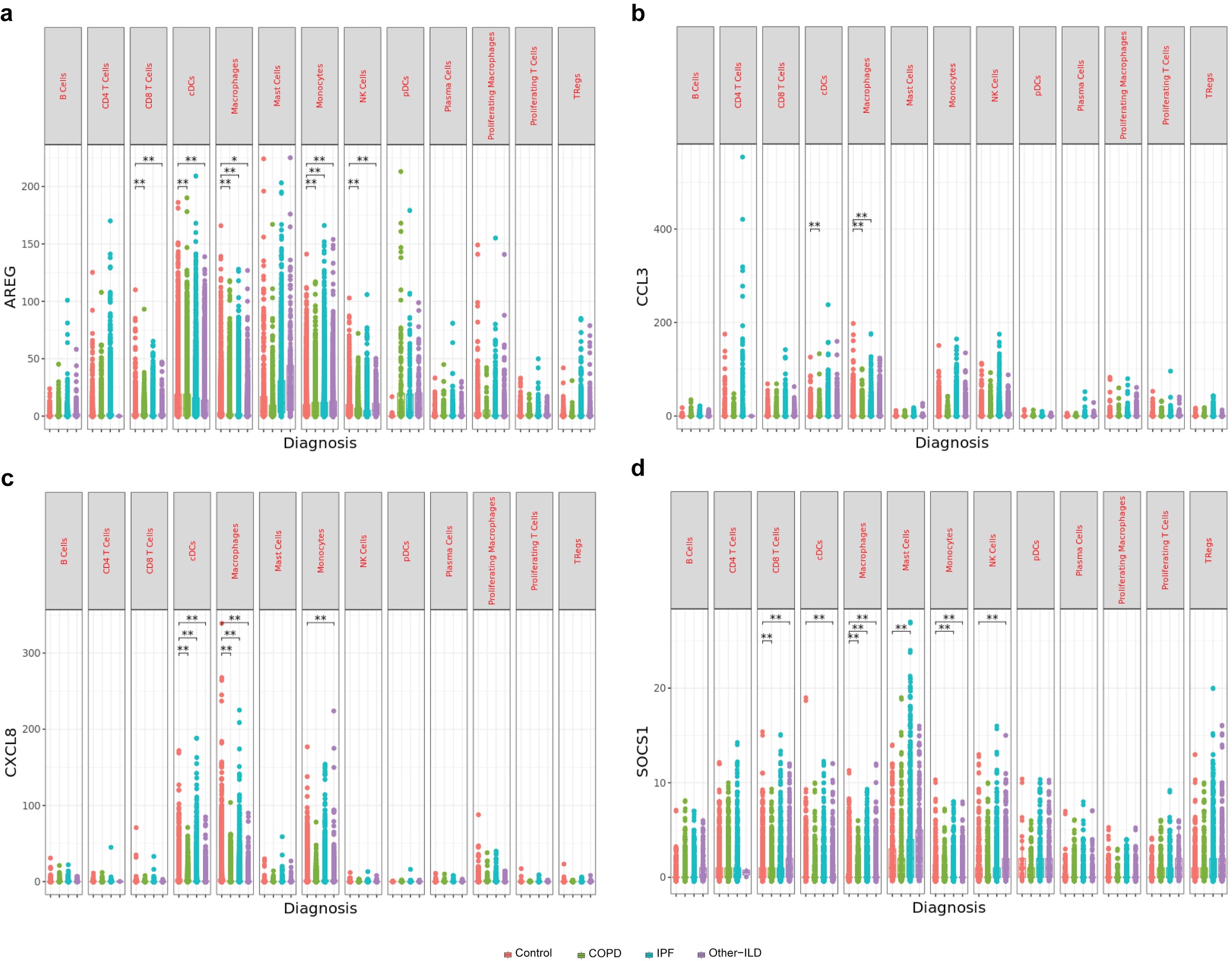

**Supplementary Fig 10**: Gene expression differences of SARS-CoV-2 immune response genes. (**a**) *AREG.* (**b**) *CCL3.* (**c**) *CXCL8.* (**d**) *SOCS1* in all immune cells. *: p_adjusted_value < 0.1, **: p_adjusted_value < 0.05, p_adjusted_value was Bonferroni adjusted from Seurat FindMarkers differential expression analysis using a negative binomial test and corrected for Age, Ethnicity, Smoking_status and Dataset.
